# Global port survey quantifies commercial shipping’s effect on biodiversity

**DOI:** 10.1101/2021.10.07.463538

**Authors:** Jose Andrés, Paul Czechowski, Erin Grey, Mandana Saebi, Kara Andres, Christopher Brown, Nitesh Chawla, James J. Corbett, Rein Brys, Phillip Cassey, Nancy Correa, Marty R. Deveney, Scott P. Egan, Joshua P. Fisher, Rian vanden Hooff, Charles R. Knapp, Sandric Chee Yew Leong, Brian J. Neilson, Esteban M. Paolucci, Michael E. Pfrender, Meredith R. Pochardt, Thomas A.A. Prowse, Steven S. Rumrill, Chris Scianni, Francisco Sylvester, Mario N. Tamburri, Thomas W. Therriault, Darren C.J. Yeo, David Lodge

**Affiliations:** Cornell University, Department of Ecology and Evolutionary Biology, 215 Tower Road, Ithaca, NY 14853, United States of America; Cornell University, Cornell Atkinson Center for Sustainability, 340 Tower Road, Ithaca, NY 14853, United States of America; University of Otago, Department of Anatomy, 270 Great King Street, Dunedin, OTA 9016, New Zealand; University of Maine, School of Biology and Ecology and Maine Center for Genetics in the Environment, 100 Murray Hall, Orono, ME 04469, United States of America; Governors State University, Division of Science, Mathematics and Technology, 1 University Parkway, University Park, IL 60484, United States of America; Golden Bear Research Center, California State University Maritime Academy. 200 Maritime Academy Drive, Vallejo, CA 94590, United States of America; University of Notre Dame, Center for Network and Data Science (CNDS), Notre Dame, IN 46556, United States of America; University of Delaware, College of Earth, Ocean, and Environment, 305 Robinson Hall, Newark, DE 19716, United States of America; Research Institute for Nature and Forest, Gaverstraat 4, Geraardsbergen 9500, Belgium; School of Biological Sciences, University of Adelaide, SA 5005, Australia; Servicio de Hidrografía Naval (Ministerio de Defensa), Av. Montes de Oca 2124, Buenos Aires, Argentina and Escuela de Ciencias del Mar, Sede Educativa Universitaria, Facultad de la Armada, UNDEF, Av. Antártida Argentina 425, Buenos Aires, Argentina; SARDI Aquatic Science and Marine Innovation SA, South Australian Research and Development Institute, 2 Hamra Avenue, West Beach SA 5024, Australia; Rice University, Department of BioSciences, 6100 Main Street, Houston, Texas, 77005, United States of America; United States Fish and Wildlife Service, Pacific Islands Fish and Wildlife Office, Honolulu, Hawaii 96850, United States of America; Oregon Department of Environmental Quality, 700 NE Multnomah Street, Portland OR 97232, United States of America; Daniel P. Haerther Center for Conservation and Research, John G. Shedd Aquarium, 1200 S. Lake Shore Dr., Chicago, IL 60605, United States of America; St. John’s Island National Marine Laboratory, Tropical Marine Science Institute, National University of Singapore, 18 Kent Ridge Road, Singapore 119227, Singapore; State of Hawaii Division of Aquatic Resources, Honolulu, Hawaii, United States of America; Museo Argentino de Ciencias Naturales “Bernardino Rivadavia”-CONICET, Av. Angel Gallardo 470, Buenos Aires C1405DJR, Argentina; Department of Biological Sciences and Environmental Change Initiative, University of Notre Dame, Notre Dame IN 14556 United States of America; M. Rose Consulting, 321 3rd Ave. Haines, AK 99827. United States of America; Marine Resources Program, Oregon Department of Fish and Wildlife, 2040 SE Marine Science Drive, Newport, OR 97365, United States of America; California State Lands Commission, Marine Invasive Species Program, 301 E. Ocean Bld., Ste. 350, Long Beach, CA 90802, United States of America; Instituto para el Estudio de la Biodiversidad de Invertebrados, Facultad de Ciencias Naturales, Universidad Nacional de Salta, Av. Bolivia 5150, A4408FVY Salta, Argentina; and Consejo Nacional de Investigaciones Científicas y Técnicas (CONICET), Salta, Argentina; Chesapeake Biological Laboratory, University of Maryland Center for Environmental Science, 146 Williams Street, Solomons, MD, United States of America; Fisheries and Oceans Canada, Pacific Biological Station, 3190 Hammond Bay Road, Nanaimo, BC, V9T 6N7, Canada; Department of Biological Sciences, National University of Singapore, 16 Science Drive, Singapore 117558, Republic of Singapore; Lee Kong Chian Natural History Museum, National University of Singapore, 2 Conservatory Drive, Singapore 117377, Republic of Singapore

## Abstract

Spread of nonindigenous organisms by shipping is one of the largest threats to coastal ecosystems. Limited monitoring and understanding of this phenomenon currently hinder development of effective prevention policies. Surveying ports in North America, South America, Europe, Southeast Asia, and Australia we explored environmental DNA community profiles evident of ship-born species spread. We found that community similarities between ports increased with the number of ship voyages, particularly if the ports had similar environments, and when indirect stepping-stone connections were considered. We also found 57 known non-indigenous taxa, some in hitherto unreported locations. We demonstrate the usefulness of eDNA-based tools for global biodiversity surveys, and highlight that shipping homogenizes biodiversity in predictable that could inform policy and management.

## Introduction

Harmful nonindigenous species transported through ballast water or biofouling of commercial ships are among the most severe threats to biodiversity and ecosystem services of marine coastal environments (Bax et al., 2003; J. Carlton, 1996; J. T. Carlton & Geller, 1993) Studies of marine traffic patterns and vessel characteristics have helped to develop more robust risks analyses and to identify global hotspots for invasion (Keller et al., 2011; Lodge et al., 2016; Seebens et al., 2013; Tidbury et al., 2016). However, lack of standardized biodiversity data at large spatial, temporal, and taxonomic scales has limited testing of this risk analyses. Cost-effective technologies suitable for quantifying species spread along individual shipping routes are required to accurately categorize shipping voyages and pathways by their risk of spreading nonindigenous species.

DNA analysis of environmental samples is an emerging method that has proven to be accurate and cost-effective for biomonitoring across a wide range of taxa and a variety of habitats (Thomsen & Willerslev, 2015) including commercial ports (Grey et al., 2018). Such analysis of environmental DNA (eDNA) also offers opportunities for quantifying global ecological impacts and evaluating risk models. Here we use a global environmental DNA metabarcoding survey to evaluate four models of ship-borne eukaryotic species spread risk. We show that eDNA analysis of port water samples, together with global data on environmental conditions and shipping traffic, enables route-specific risk analyses of ship-borne species spread (Porter & Hajibabaei, 2018; Saebi, Xu, Kaplan, et al., 2020; Thomsen & Willerslev, 2015; Xu et al., 2016). By implementing a standardized and efficient eDNA port survey method (Grey et al., 2018) we identified biological similarities between 19 ports and related them to four different risk estimates of ship-mediated species transport, while concurrently considering the effects of environmental similarity (Keller et al., 2011) and biogeography (Costello et al., 2017) on global community compositions. Extending this approach to further ports would allow for evaluation and refinement of risk analyses that could better inform policy and management of ship-born nonindigenous species spread.

Our four alternative shipping-risk estimates derived from long-term ship traffic data consider two different aspects of the species spread risk along routes in the global shipping network. The first aspect considered route spread risk as either function of the number of ships traveling between two ports (“traffic risk”) or as a function describing the amount of ballast water (“ballast risk”) transported between ports, based on correlations between ballast discharge volume and probability with ship size, ship type, and voyage time (Saebi, Xu, Grey, et al., 2020; Saebi, Xu, Kaplan, et al., 2020). The second aspect considered the importance of either direct port-to-port voyages (“direct connection risk”) or indirect connections through intermediary ports (“stepping-stone connection risk”). Stepping-stone spread occurs when species are transported from the origin port to an intermediary port, and then picked up by another ship that transports them to the destination port. In this scenario, it is possible to have species spread between two ports even without direct ship traffic between them. While stepping-stone transport of nonindigenous species has been documented at regional scales (Apte et al., 2000), its relative importance at the global scale is unknown.

To evaluate our risk models, we related eDNA-derived biological community profiles, estimates of shipping risk, environmental similarity and shared biogeography in linear models that also considered all two-way interactions of these factors, and random effects related to each port-to-port shipping connection. We then used backward AIC selection to identify and rank the best model for each risk metric. We expected increased shipping (expressed through shipping-risk estimates) to increase biological similarities between connected ports, while keeping in mind that environmentally similar ports and ports within the same biogeographic region would be biologically similar also in the absence of shipping, thereby providing a backdrop to ship-mediated species spread.

## Results

Nineteen of 21 port that we sampled yielded high quality sequence data (Fig. 1a, Table S1). Port locations spanned seven of the 30 marine biogeographic realms defined by Costello et al. (Costello et al., 2017), with the majority situated in the North Pacific, Caribbean and Gulf of Mexico, and Northeast Atlantic realms (Figure 1a). In total we obtained 11,975 eukaryotic Amplicon Sequence Variants (ASVs, (Callahan et al., 2017)) representing both planktonic and benthic species (Fig. S1). Port-to-port Unifrac distances (Lozupone et al., 2011), which are phylogenetic measures of community dissimilarities, ranged from 0.49 to 0.94 (mean: 0.79 ± 0.07) (Figure 1b). Port-to-port values of our four shipping risk estimates were similar each other, but dissimilar to the patterns observed for port-to-port environmental distances (Figure 1b).

**Fig. 1.**
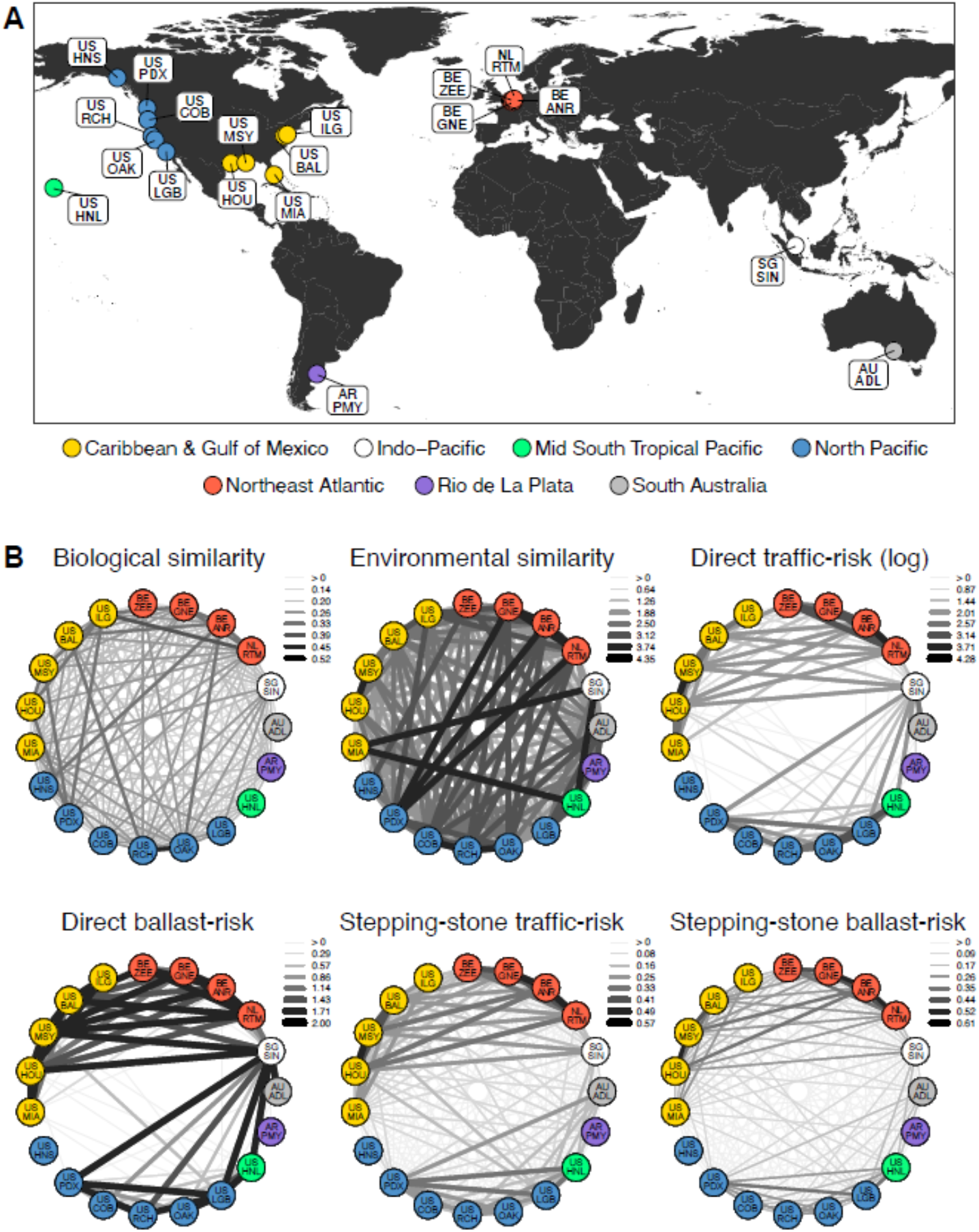
Sampling extent and variable summary. **(A)** Sampled ports, colored by biogeographic realms (Costello et al., 2017), included Adelaide (AU-ADL), Antwerp (BE-ANR), Baltimore (US-BAL), Coos Bay (US-COB), Ghent (BE-GNE), Haines (US-HNS), Honolulu (US-HNL), Houston (US-HOU), Long Beach (US-LGB), Miami (US-MIA), New Orleans (US-MSY), Oakland (US-OAK), Portland (US-PDX), Puerto Madryn (AR-PMY), Richmond (US-RCH), Rotterdam (NL-RTM), Singapore (SG-SIN), Wilmington (US-ILG), and Zeebrugge (BE-ZEE) **(B)** Summaries of network-like relationships between ports expressing biological similarity, environmental similarity, and different shipping-risk estimates. To allow for direct comparison, we inverted biological and environmental dissimilarities by subtracting each dissimilarity value from either the theoretical (1 for Unifrac distance) or observed (4.36 for environmental dissimilarity) maximum. Line color and width are scaled according to each metric, where thick black lines represent the highest values and thin light-grey lines the lowest values.

Regression modeling results supports our prediction that shipping effect on biodiversity between ports can be detected amongst a backdrop of environmental similarity and, to a lesser extent, biogeographic patterns. All AIC-selected models identified significant effects of environmental similarity (temperature and salinity (Keller et al., 2011)) on biological similarity (Unifrac distance) between ports (Table 1) and connected port pairs with similar environments tended to have smaller Unifrac distances (i.e. more similar biological communities; Fig. S2). Furthermore, ports within the same biogeographical realm (Costello et al., 2017) had the most similar biological communities (Fig. S3) although this effect is only statistically significant in the direct risk models.

**Table 1.** Model summaries. Selected linear models including environmental similarity (ENV), traffic risk (SHP), and biogeographic realms (BGR). Ports of origin (ORG) and destination (DEST) were modelled as random factors. AIC: Akaike Information Criteria.

| SHIP RISK METRIC | AIC-SELECTED MODEL | AIC |
| --- | --- | --- |
| Stepping-stone traffic risk | UNIFRAC = ENV+ SHP + ENV*SHP + (ORG) + (DEST) | -<br>479.34 |
| Stepping-stone ballast risk | UNIFRAC = ENV + SHP + ENV*SHP+ (ORG) + (DEST) | -<br>478.78 |
| Direct ballast risk | UNIFRAC = ENV + BGR+ SHP + BGR*ENV+BGR*SHP + (ORG) + (DEST) | -<br>478.51 |
| Direct traffic risk | UNIFRAC = ENV + BGR + ENV*BGR+ (ORG) + (DEST) | -476.9 |

Consistent with our predictions, shipping increased the biological similarity between ports. All but one of the AIC-selected models included shipping as an explanatory variable (Table 1). Contrary to our predictions, models considering traffic risk and ballast risk had similar AIC values, suggesting that considering the complexities of ballast discharge beyond the number of voyages added little explanatory power. Models considering stepping-stone connection risk ranked higher than models considering direct connection risks, highlighting the potential need to consider indirect spread through intermediary ports when estimating route-based spread risks.

Most informative to explain species spread along shipping routes was a relatively simple model considering only environmental similarity, stepping-stone traffic connections between ports, and interactions of these variables (Table 1, first row, with lowest AIC). Further exploration of this model revealed that shipping risk increased biological community similarity, particularly on routes whose origin and destination are environmentally similar (Fig. 2). This observation is consistent with the expected pattern of ship-borne species spread, as organisms are unlikely to survive in vastly different environments, regardless of how many times they are introduced.

**Fig. 2.**
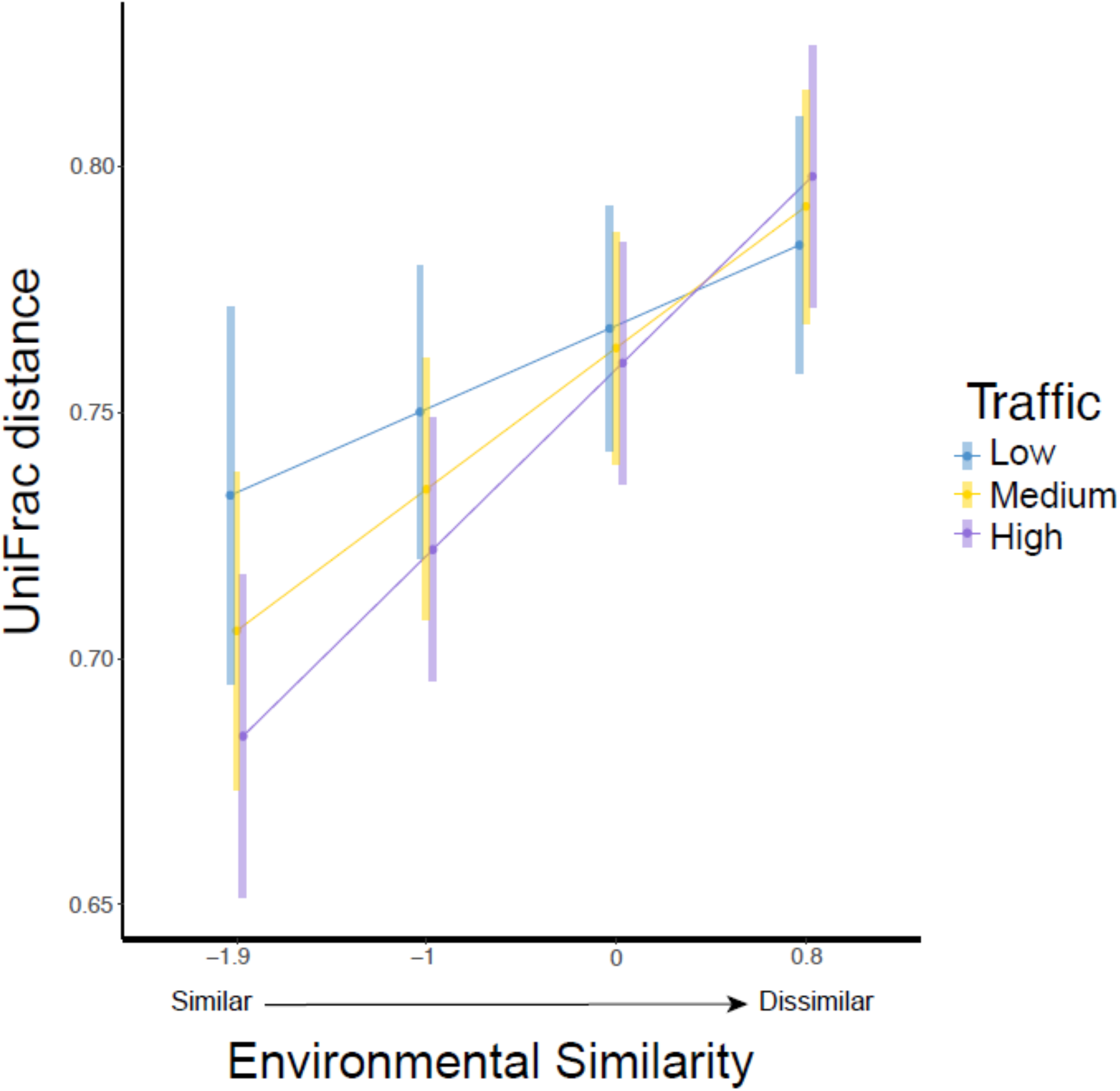
Observed Relationship between biological similarity, environmental similarity, and stepping-stone connection risk. Biological similarity between ports is best explained by the interaction of stepping-stone vessel traffic and the environmental similarity between the ports of origin and destination (Table 1). Medium = Mean; Low = Mean - 1SD; High = Mean + 1SD. Bars represent the ± 95% CI.

Overall, our analysis revealed that shipping homogenizes biodiversity among ports when evaluated against a backdrop of high eukaryotic biodiversity shaped primarily by environmental similarity. Across all models, environmental similarity (expressed through temperature and salinity) had the largest effect on UniFrac distances, as we found coefficients associated with shipping risk variables to be much smaller than those associated with environmental similarity. The smallest coefficients were those of the biogeographic variables (Table S2). Our modelling results are robust, as analogous analyses adding two more ports at a shallower sequencing depth yielded similar results, except with biogeographic realms emerging as an additional explanatory variable (Table S3).

A strength of our analyses is the incorporation of taxon-agnostic eDNA profiles from a vast range of taxa from many samples, thereby including information from hitherto undescribed or cryptic species, and overcoming limitations of eDNA studies that attempt low-level taxonomic classification using binomial nomenclature. Nonetheless, species identification with binomial nomenclature is still a critical component of invasion risk management. Accordingly, to further explore overlapping distribution of species among sampled ports, we focused on the subset of ASVs with taxonomic annotations matching nominal species as included in the World Register of Introduced Marine Species (WRiMS) (Ahyong et al., n.d.).

We identified 273 ASVs of which 79 (28.9%), matched equally well to more than one nominal species, indicating possible reference database errors or a lack of taxonomic resolution. The remaining 194 taxonomically unambiguous ASVs represented 11 classes and 57 unique nominal species (Fig. 3), with the solitary tunicate *Ciona savignyi* (41 ASVs) and reef-forming polychaete *Ficopomatus enigmaticus* (42 ASVs) comprising over 40% of all reads for these ASVs. Two of these 57 species were found in non-native areas where they had not been previously reported (Table S4) - *Pseudocalanus elongatus* (Copepoda) at Coos Bay, and *Botrylloides leachii* (Ascidiacea) at Richmond and Oakland. One species, *Oithona davisae*, was found slightly outside of its reported native range in Singapore. All other species were found within their previously reported ranges.

**Fig. 3.**
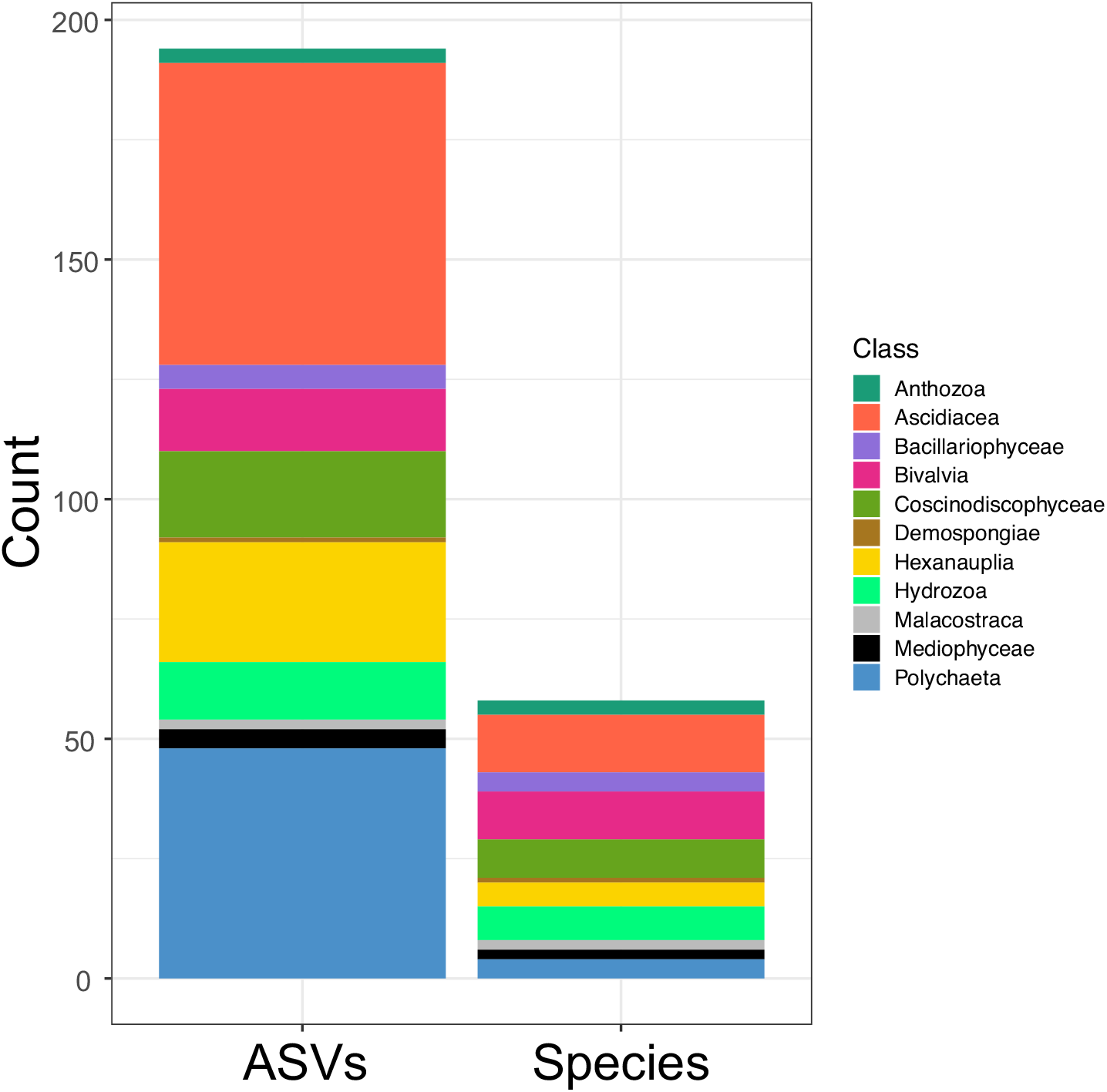
Evaluation of marine nonindigenous species. Count of ASVs and number of unambiguous nominal species (99% identity) matching to the WRiMS database of marine invasive species colored by taxonomic Class.

## Discussion

Our work contextualizes previous, more spatially limited studies on nonindigenous and invasive taxa in ports, including traditional and eDNA approaches (Borrell et al., 2017; Verna et al., 2016). We provide *in situ* evidence of the relative effects of shipping and the environment on biological homogenization at the global scale. We also reveal the importance of stepping-stone connections for species spread through the global shipping network, which highlights the need to consider the whole shipping network to predict species spread (Saebi, Xu, Kaplan, et al., 2020; Xu et al., 2016). While it is generally accepted that differences among ship types result in unequal risks of species transfer (Davidson et al., 2018; Drake & Lodge, 2004), including ship and voyage parameters influencing ballast discharge in our models had no significant effect on our ability to explain the biological similarities between ports. Although counterintuitive at first glance, it is possible that ship-borne species spread from biofouling or other vectors such as aquaculture obscures the ballast discharge effect in the studied ports (Ojaveer et al., 2018). Evaluation of models with more detailed shipping-risk estimates including biofouling-specific parameters such as vessel speeds and port residence times are needed to determine if this is the case.

Collectively, our results confirm that species spread risks differ among ports in predictable ways, and that relatively simple models yield informative predictions for invasion management. Our most informative model used a shipping risk estimate based on voyage counts between ports, such counts are routinely collected and readily available. The observed significant interaction between shipping and environment suggests that high volume shipping routes connecting environmentally similar ports are the highest risk routes, especially when passing through ports serving as hubs to many other ports, such as, but not limited to, Singapore. Prioritizing these risky routes with targeted management or biomonitoring would, in our opinion, be a more efficient way to reduce global ship-borne species spread.

The lack of a consistently strong signal of biogeographic realms in our results is puzzling, and could be caused by current biogeographic delineations being biased either taxonomically, spatially, or temporally and/or by ports being distinct habitats that are generally quite similar around the world (Bulleri & Airoldi, 2005; Forrest et al., 2013; Lambert & Lambert, 2003; Megina et al., 2016; Piola & Johnston, 2008). It could also be that the limited temporal scope of our study (one sampling event per point) does not reflect β-diversity patterns over longer time-scales (e.g., months to years). Further investigations including ports from more bioregions and other coastal habitats sampled throughout the year are needed to explore such non-exclusive hypotheses.

We demonstrate that eDNA analysis is useful for biomonitoring at a global scale (Campbell et al., 2007). Environmental DNA analysis yields robust β-diversity estimates suitable for hypotheses testing related to the many factors that influence observed biodiversity patterns. For species monitoring, the current drawbacks of eDNA applications are their inaccuracy in assigning taxonomy to some sequences when reference data is deficient, and their susceptibility to biased abundance estimates (Balvočiūtė & Huson, 2017; Kelly et al., 2019). Regardless, we discovered sequence variants highly similar to many known nonindigenous species and derived important insight on their global distributions, which could be further explored with colour-based DNA indicator reactions (Notomi et al., 2015) or enrichment (Wilcox et al., 2018) targeting specific species. Likewise, further exploitation of eDNA techniques may soon enable population genetic inquiries (Sigsgaard et al., 2017) and possibly, abundance estimates (Spear et al., 2021), also in across global ports.

Overall, our work suggests that route-specific management may be a feasible and more effective approach to manage ship-based species invasion risks than then current global “one-size-fits-all” management (Molnar et al., 2008; Wang et al., 2020). While the effect of biogeography and the most appropriate shipping risk metric remain uncertain, shipping clearly homogenizes biodiversity in predictable ways. We show that relatively simply obtained and readily available shipping risk and environmental similarity variables can be used to prioritize route-based management. Future port eDNA surveys including other bioregions and ports experiencing changes in environment and shipping could further refine understanding of ship-born species spread and empower efficient policy development.

## Materials and Methods

### Water sample collection

We collected and analyzed five water samples in each of 21 ports across seven biogeographic realms (3, 7, 11, 13, 17, 24, and 26 as defined in Costello *et al*. (Costello et al., 2017) including Australia, China, Oceania, Europe and the Americas (Fig.1). To control for diurnal, tidal, and seasonal variation, all samples were collected during daylight hours at slack hightide after at least a 12-hour period of no rain during the summer or early fall (June-November) (Table S4). Sampling followed a protocol developed specifically for ports and published in Grey *et al*. (Grey et al., 2018). Briefly, each sample consisted of 250 mL surface water, which was stored at 4°C for a maximum of 12 hours after collection. Within that time, water was filtered through a 0.45 μm cellulose-nitrate membrane, immediately immersed in 700 μL Longmire’s buffer and stored at room temperature for up to two weeks, and then at -20°C until extraction. If 250 mL could not be filtered through a single membrane, as was the case at some ports with high turbidity and algae, then additional membranes were used and stored in the same microtube. For each site, a collection “blank” sample was taken by filtering 250 mL of bottled or tap water, which was subsequently treated identically to other samples.

### eDNA extraction

DNA extractions for the samples and collection blanks (see below) taken at the ports of Adelaide, Chicago, and Singapore were done using phenol-chloroform in a dedicated PCR-free lab at the University of Notre Dame as described by Grey *et al*. (Grey et al., 2018). Total DNA from all other samples and collection blanks were extracted in an eDNA facility at Cornell University (UV light for 8 h d^-1^, HEPA filtered air under positive pressure, personnel wearing full body suits, face-shields, and breathing masks) using a Blood and Tissue DNA extraction kit (Qiagen, Hilden, DE-NW) and an eDNA optimized protocol (Spens et al., 2017).

### Library design, amplification and sequencing

18S rRNA amplicon libraries were prepared using two primers targeting the eukaryotic V4 region (18S_574F and 18S_952R) that were previously evaluated by Hadziavdic et al. (Hadziavdic et al., 2014). For 19 ports (all but Adelaide, Chicago, and Singapore), we followed a single-step amplification approach using a single set of primers including Illumina adapters and a 12-bp barcode unique to each sample (Fig. S4).

Threefold PCR amplifications were carried out for all samples to account for amplification bias. Each reaction of 25μL total volume contained 2μL template DNA, 1.5 mM MgCl_2_, 1× Colourless GoTaq Flexi Buffer (Promega, Fitchburg, US-WI), 0.25 mM of each dNTP, 0.2 μM of forward and reverse primer and 1.25 units GoTaq Hot Start DNA Polymerase (Promega). After initial denaturation at 95°C for 2:00 min, we performed 40 amplification cycles (95°C for 45 s, 49°C for 1 min, 72°C for 1 min), and a final elongation at 72°C for 5 min. Amplicon triplicates were pooled, visualized on 2% agarose gels, and then purified using Mag-Bind Total Pure NGS beads (Omega Bio-Tek, Norcross, US-GA). DNA concentration for all samples was quantified using PicoGreen reagent (Thermo Fisher Scientific, Waltham, US-MA) and a Spectramax M2 multi-detection microplate reader (Molecular Devices, San Jose, US-CA). Libraries were prepared by combining samples in equimolar ratios, and pair-sequenced (2 × 250) on an Illumina MiSeq platform (Fig. S5). Libraries for the ports of Adelaide, Chicago, and Singapore were previously generated by some of the authors using the protocols described in (Grey et al., 2018).

### Blanks and Positive controls

Alongside port water DNA libraries, we generated libraries for three different blank types (collection, extraction, and amplification controls) and three positive controls: One containing genomic DNA of *Danio rerio*, and two equimolar mixtures of either 4 freshwater fishes (*Notropis topeka, Noturus taylori, Umbra limi, Thoburnia atripinnis*) or 6 marine fishes (*Pseudanthias dispar, Ecsenius bicolor, Macropharyngodon negrosensis, Centropyge bispinosa, Salarius fasciatus, Amphiprion ocellary*; Fig. S5). Reference sequences for all of the above species were generated using genomic DNA and Sanger sequencing on an ABI 3730xl. Sequencing failed for *Macropharyngodon negrosensis*.

### Sequence data processing

Amplicon Sequence Variants (ASVs) were generated from combined 18S amplicon sequence data with Qiime v2 2019-04 (Bolyen et al., 2019), after using BBtools to include sequence data from (Grey et al., 2018) into this work. To do so, reads were initially imported, paired, and filtered using Qiime’s import functions with default parameters. Read pairs were then subjected an additional round of adapter trimming using Cutadapt v1.18 (Martin, 2011), denoised using DADA2 (v1.6.0)(Callahan et al., 2016) and an Expected Error value of 9 (Edgar & Flyvbjerg, 2015).(Edgar & Flyvbjerg, 2015) Prior to merging, reads were trimmed to 220 nucleotides. The resulting unique Amplicon Sequence Variants (ASVs) were then filtered using Qiime’s taxonomic classifier Vsearch (Rognes et al., 2016) to obtain eukaryotic ASVs as follows: After appending our positive control reference sequences to the Qiime release of the SILVA database v132 (Pruesse et al., 2007), we first ran a separate *in silico* experiment to optimize the assignment settings of Vsearch. Specifically, we optimized the classifier to unequivocally identify the species of the positive controls and to yield maximum correspondence to the analogous taxonomy assignments obtained using the Blast algorithm (Camacho et al., 2009) and the nt/nr NCBI database (Benson et al., 2018). We then used the optimized Vsearch classifier and the SILVA database v132 to identify and retain all Eukaryotic ASVs (*-p-per-identity*= 0.875, *-p-min-consensus*= 0.5, -*p-query-cov*= 0.9).

To account for contamination in the field and lab, we inspected and subtracted ASVs present in the negative controls (see blanks indicated in Fig. S5) from all subsequent datasets. Sequence run read outputs and read counts at each bioinformatics stage are shown in Fig. S6.

### Eukaryotic ASV rarefaction and port sampling effort

To account for different read depths across samples, we generated rarefaction curves using Qiime to determine the minimum number of ASVs describing the eukaryotic diversity found among the sampled ports. To do so, we rarefied all samples at depths between 1 to 55,000 sequences in steps of 1,000, repeating each step 4 times. Based on the resulting accumulation curves we initially determined 49,900 ASVs/port sample to be sufficient. Because choosing specific plateauing values is to some extent arbitrary, we also consider a second, lower, value of 37,900, thereby allowing inspecting of two more ports (Fig. S7).

For each of these two rarefied datasets, we then used a bootstrapping approach to determine the minimum number of samples per port that result in stable estimates of Unifrac distances between port pairs. Our bootstrapping algorithm initially divided the full Unifrac distance matrix (including all samples and ports) into unique port-pair matrices. From each of these port matrices, we then generated a series of bootstrapped squared, symmetric matrices of varying dimensions ranging from 1×1 to 7×7 samples/port (n=10,000/dimension). For each bootstrapped matrix a mean distance value was calculated. Finally, the median absolute deviation (MAD for each port pair and sampling dimension) was calculated and plotted (Fig. S8 and S9). Based on the pattern of MAD decline, we determined that, for both rarefied datasets, 5 samples were sufficient to provide robust estimates of Unifrac distances between port pairs.

Given the above results, we generated two different datasets by randomly selecting five sufficiently covered samples per port and rarefying them to a depth of either 49,900 or 37,900 eukaryotic ASVs. The former dataset consists of 10,279,400 and excludes the ports of Nanaimo and Vancouver while the latter one comprises 8,262,200 eukaryotic ASVs and included all sampled ports.

### Estimation of Unifrac distances between port pairs

For each of the two rarefied datasets, we calculated the Unifrac distances between ports using Qiime. To do so, we first obtained reference trees by aligning eukaryote ASV’s using MAFFT (Katoh & Standley, 2013) with automatic algorithm and parameter selection. This initial alignment was then post-processed to retain only consenting columns supported by at least 50% of ASV’s and allowing only columns with < 10% gaps across the entire alignment. Finally, reference trees were obtained using Fasttree (Price et al., 2010) with the default settings, and unweighted Unifrac distances were calculated among all port pairs.

### Modelling of traffic- and ballast-associated spread risks

NIS (non-indigenous species) spread via shipping activity depends on multiple environmental and geographical factors, as well as complex interactions in the global shipping network. For example, many ships do not discharge their entire ballast water in their first port of call. As a result, the remaining ballast water discharged by the ship at the intermediate stops poses the risk of NIS transfer from the original port to the intermediate stops. To account for more such patterns of NIS spread, we followed the approach of (Saebi, Xu, Grey, et al., 2020; Saebi, Xu, Kaplan, et al., 2020) to obtain 4 different traffic- and ballast-associated risks. We used eight years of shipping data from 1997 to 2018 obtained from Lloyd’s List Intelligence, an Informa Group Company (LLI, New York, NY, USA). We calculated the risk of spread based either on direct traffic, stepping-stone traffic, direct ballast, or stepping-stone ballast. In the two ballast network models, the spread risk was obtained by first calculating the introduction probability for all ships travelling from port *i* to port *j*. For each ship *v* travelling during time Δ*t*_*ij*_^*v*^ we applied:

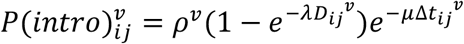

Where *D*_*ij*_^*v*^, ρ^v^, ƛ are the amount of ballast water discharged at the destination, the efficacy of ballast water management for the route, and the species introduction potential per volume of discharge.

In the stepping-stone network models, for each port pair, we included an indirect risk of spread by calculating a Jaccard coefficient by dividing the number of ports directly connected to both ports by the total number of direct connections to either port in the pair.

### Statistical modelling

A critical first step in understanding the role of shipping traffic in the homogenization of marine biota is the selection of an appropriate set of models covering reasonable hypotheses. Marine species distributions are known to be influenced by temperature, salinity, and biogeography (e.g. (Costello et al., 2017; Keller et al., 2011)). Therefore, we considered the additive model:

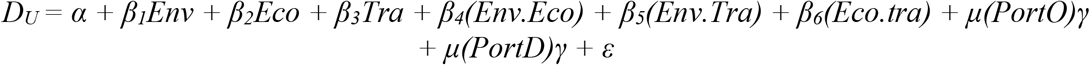

Where *D*_*U*_ is the Unifrac distance between port pairs, the ports of origin (*PortO*) and destination (*PortD*) are random effects, shipping traffic (*Tra*) and environmental similarity (*Env*) are fixed factors, and the biogeographic distance between port pairs (*Eco)* is a binary fixed effect that reflects if the ports of origin and destination are in the same marine realm as defined by Costello *et al*. (Costello et al., 2017) or not. We ran 4 different models using voyage frequency, with or without consideration of “stepping-stone connection risk” estimates, which were calculated as Jaccard indices for each port pair. The environmental similarity between ports was estimated as described elsewhere (Keller et al., 2011). These complex models consider all additive effects and the potential effect of all the two-way interactions. To avoid over- or under-fitting, we then used a backward AIC variable selection approach. Statistical modeling and model selection were done using centered and scaled response variables and the *lme4* package (Bates et al., 2015) with the R programming language (R Core Development Team, 2019).

### Marine invasive species identification

Taxonomic assignment of ASVs was performed using the Blast algorithm version 2.9 (Camacho et al., 2009), the April 2019 NCBI nr/nt reference database (Benson et al., 2018) (excluding subject sequences solely identified as “environmental sample”), and a minimum e-value of 10^−5^. For each query-subject pair, we kept the top 5 high-scoring alignments (>98.5% percent identity) and excluded “ambiguous” assignments where more than one species matched equally well to the query sequence. Unambiguously identified species were then matched to the World Register of Introduced Marine Species, WRiMS (Ahyong et al., n.d.) and the species ports’ presence data was contrasted with the known distributions of these species according to the World Register of Marine Species, WoRMS (WoRMS Editorial Board, 2019) (Table S2). R packages *phyloseq* (McMurdie & Holmes, 2013) and *tidyverse* (Wickham, 2017) were used to import and ASV data and Blast results. We used *taxonomizr* to obtain taxonomic information of the best Blast hits.

## Acknowledgments

For helpful discussions and/or logistic collaboration we thank Kimberly Howland, Anna Lacoursiere, Yiyuan Li, and Aibin Zhan, Demetrio Boltovskoy, Pedro Brandão, Juul Lorre, Dias F. Pinto, Kristy Deiner, Vanessa Hodes, Jill Remeysen, Lindsay Schaffner, Evangelina Schwindt, Peter Avis, and Giles Hooker.

## Author contributions

Conceptualization: EKG, MP, DML

Methodology: PC, JA, EKG, DML, MS

Investigation: PC, JA, KA, EKG, DML, MS

Visualization: JA, KA, PC, EKG

Supervision: JA, DML, NC

Writing—original draft: PC, JA, EKG, DML

Writing—review & editing: all authors

## Data availability

Further details on data and analyses are included in the Supplementary Material. All data, code, and materials used in the analyses will be available electronically via Zenodo [link TBD]. Sequences will be available through the NCBI Sequence Read Archive [Project # and link TBD]

