## Supplemental Information for "Global port survey quantifies commercial shipping’s effect on biodiversity"

**This supplement includes:**

Figs. S1 to S7.

S1: NCBI-based taxonomy of ASVs by port.
S2: Relationship between environmental and Unifrac distances.
S3: Relationship between biogeographic realm sharing and Unifrac distances.
S4: Schematic representation of the library primers.
S5: Schematic representation of sequencing runs and libraries
S6: Read counts of each sample across bioinformatic stages
S7: ASV read accumulation curves by port.
S8: Pattern of Unifrac MAD decay with increasing sampling effort (shallow rarefied dataset).
S9: Pattern of Unifrac MAD decay with increasing sampling effort (deep rarefied dataset).

Table S1 to S4.

S1: Sample collection dates and coordinates.
S2: Coefficients of the AIC-selected models (deep rarefaction).
S3: AIC-selected models: deep and shallow rarefied datasets.
S4: Non-indigenous species sequence counts by port.

**Fig. S1.**

ASVs by eukaryotic phylum and port. Phyla designations followed NCBI taxonomy. NA represents ASVs that could not be assigned to a eukaryotic phylum. Port abbreviations: AD – Adelaide, AW – Antwerp, BT – Baltimore, CB – Coos-Bay, GH – Ghent, HN – Honolulu, HN – Haines, HT – Houston, LB – Long Beach, MI – Miami, NA – Nanaimo, NO – New-Orleans, OK – Oakland, PL – Portland, PM – Puerto Madryn, RT – Richmond, RT – Rotterdam, SI – Singapore, VN – Vancouver, WL – Wilmington, ZB – Zeebrugge.

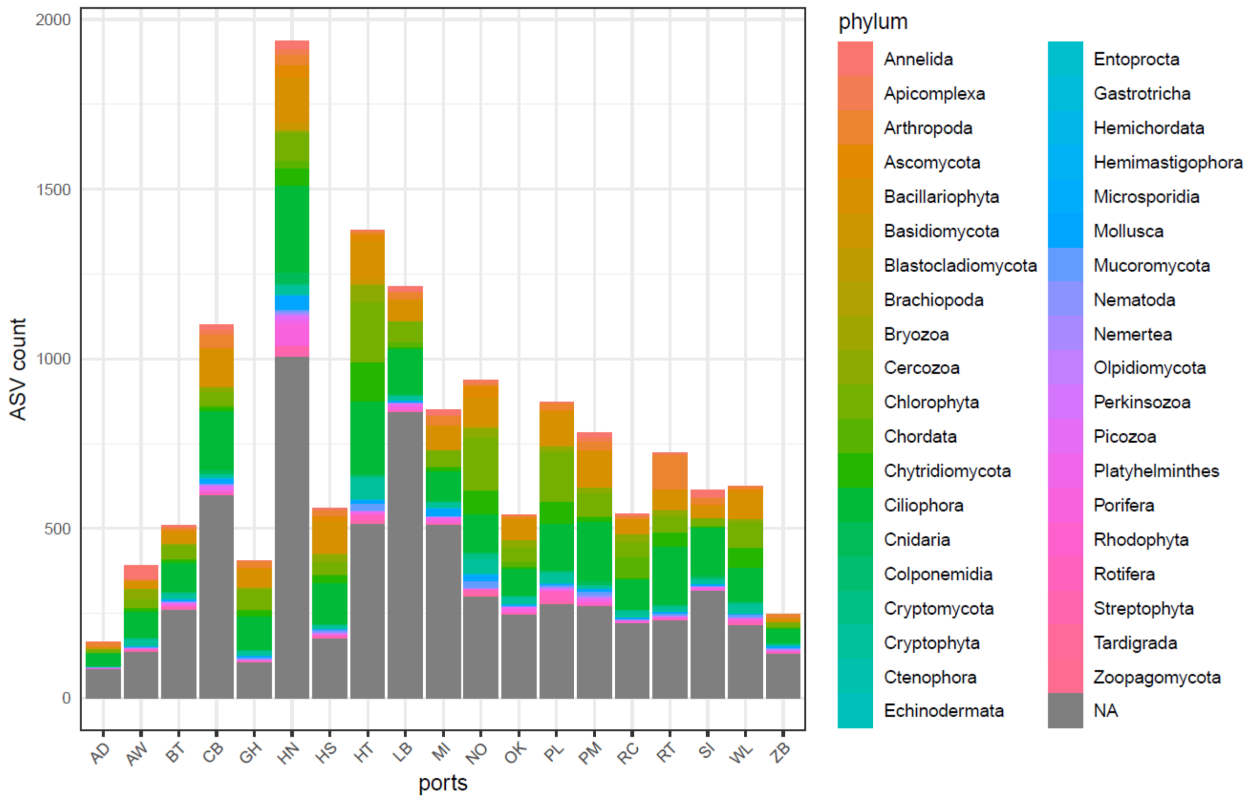

**Fig. S2.**

Relationship between environmental distances and Unifrac dissimilarities between ports. Black dots represent observed values, red dotted line represents simple linear correlation between the two variables, and blue line and grey shading represents smooth conditional means fit plus or minus standard error using the R *tidyverse* geom_smooth() function with default settings.

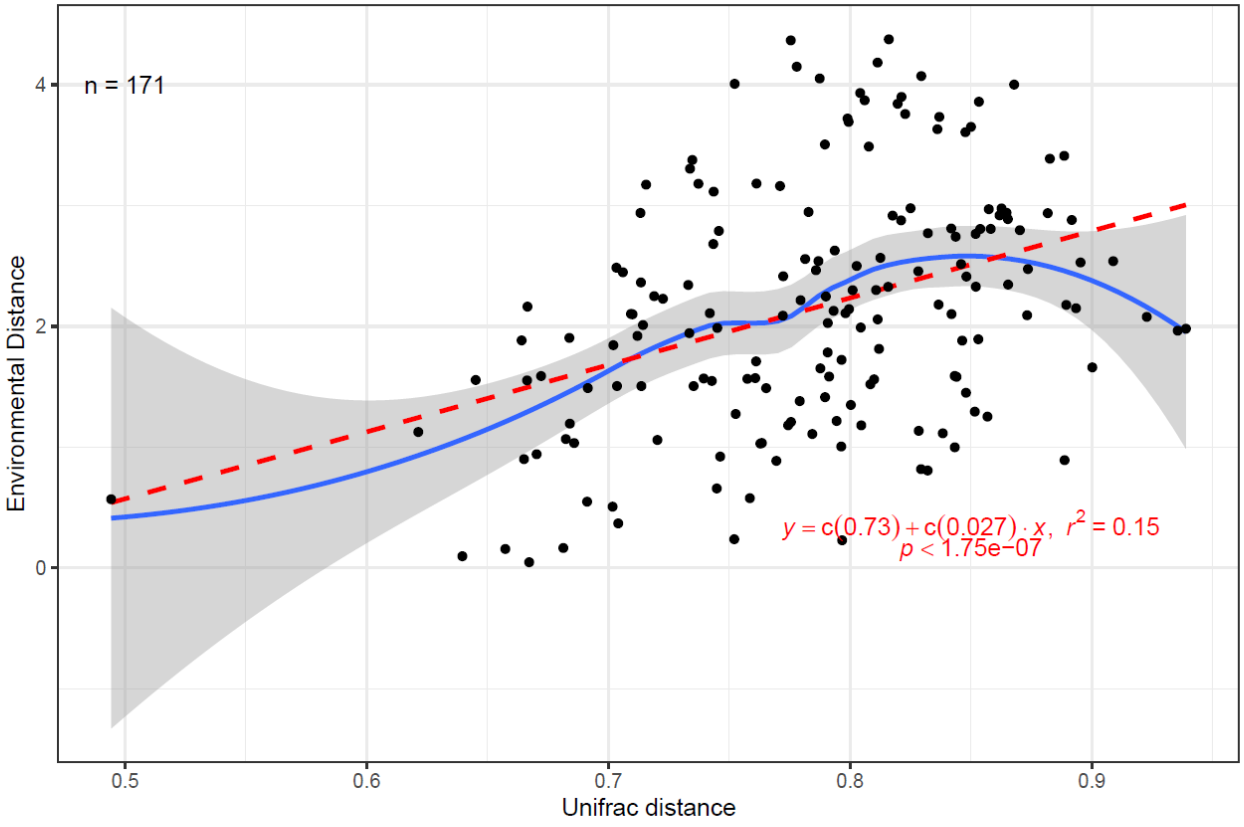

**Fig. S3.**

Relationship between biogeographic realm sharing (pink=not shared, blue=shared) and Unifrac distances between ports.

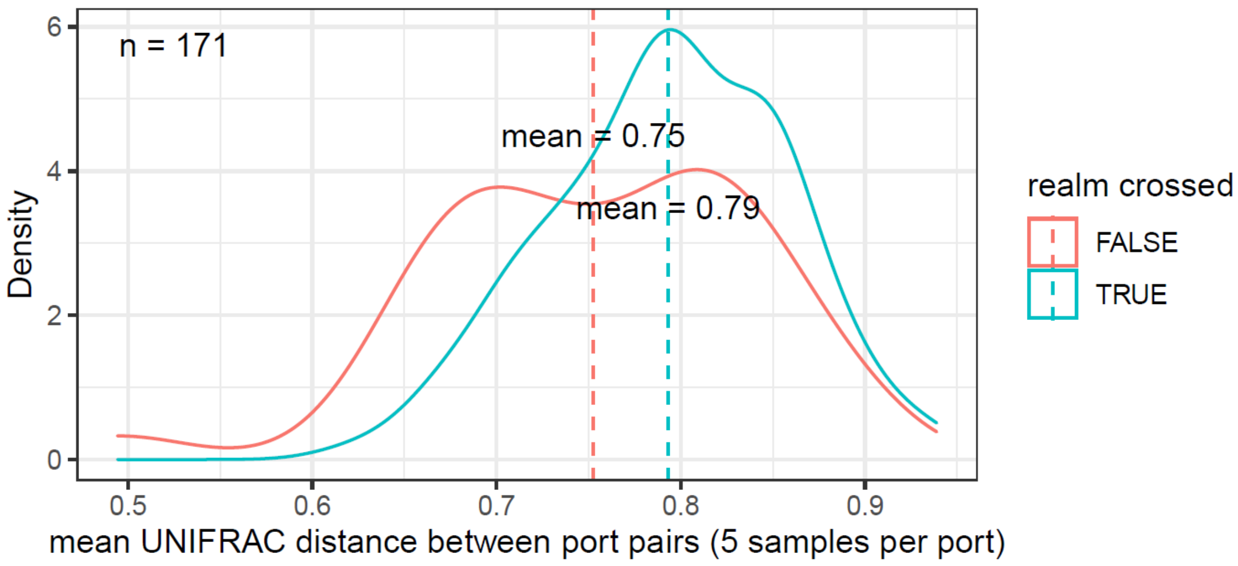

**Fig. S4.**

Schematic representation of the library primers.

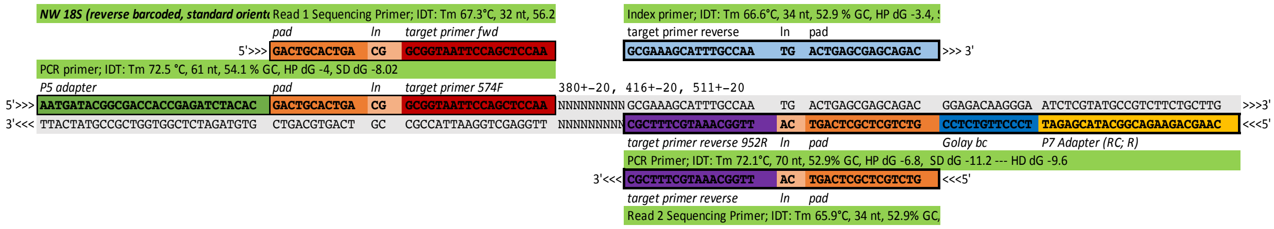

**Figure S5**

Schematic representation of sequencing runs and libraries including blanks and positive controls that were conducted at Cornell University for this study including (a) test run with Pearl Harbor samples, (b) run 1 with samples and blanks from 12 ports and mock communities, (c) run 2 with samples and blanks from 12 ports and mock communities and (d) run 3 with samples and blanks from 4 ports and mock communities. For all figures. “DW” = collection blank, “LM = Longmire’s buffer blank, “NG” = PCR negative blank, “XB” = extraction blank, “fw_mock” = freshwater mock community, “sw_mock” = saltwater mock community, and “zebra” = zebra mussel *Dreissena bugensis*. For details on the sequencing runs for the Adelaide and Singapore samples please see Grey et al. 2018 (*6*).

a. Cornell Test Run

| | 1 | 2 | 3 | 4 | 5 | 6 | 7 | 8 | 9 |
|---|---|---|---|---|---|---|---|---|---|
| A | *_MSq_0017 | *_MSq_0018 | *_MSq_0019 | *_MSq_0020 | *_MSq_0021 | *_MSq_0022 | *_MSq_0023 | *_MSq_0024 | *_MSq_0025 |
| | PH1 | PH2 | PH3 | PH4 | PH5 | PH6 | PH7 | PH8 | H20 |
| B | *_MSq_0029 | *_MSq_0030 | *_MSq_0031 | *_MSq_0032 | *_MSq_0033 | *_MSq_0034 | *_MSq_0035 | *_MSq_0036 | *_MSq_0026 |
| | PH9 | PH10 | PH11 | PH12 | PH13 | PH14 | PH15 | PH16 (xblk) | H20 |

b. Cornell Run 1

| | Rotterdam | Puerto Madryn | Buenos Aires | Coos Bay | Richmond | Haines | Oakland | New Orleans | Nanaimo | Vancouver | Portland | Antwerp | |
|---|---|---|---|---|---|---|---|---|---|---|---|---|---|
| | 1 | 2 | 3 | 4 | 5 | 6 | 7 | 8 | 9 | 10 | 11 | 12 | |
| A | RT-01 | PM-01 | BA-01 | CB-01 | RC-01 | HS-01 | OK-01 | NO-01 | NA-01 | VN-01 | PL-01 | AW-01 | sample |
| | 18S952R_MiSeq_0113 | 18S952R_MiSeq_0114 | 18S952R_MiSeq_0115 | 18S952R_MiSeq_0116 | 18S952R_MiSeq_0117 | 18S952R_MiSeq_0118 | 18S952R_MiSeq_0119 | 18S952R_MiSeq_0120 | 18S952R_MiSeq_0121 | 18S952R_MiSeq_0122 | 18S952R_MiSeq_0123 | 18S952R_MiSeq_0124 | barcode |
| B | RT-02 | PM-02 | BA-02 | CB-02 | RC-02 | HS-02 | OK-02 | NO-02 | NA-02 | VN-02 | PL-02 | AW-02 | |
| | 18S952R_MiSeq_0125 | 18S952R_MiSeq_0126 | 18S952R_MiSeq_0127 | 18S952R_MiSeq_0128 | 18S952R_MiSeq_0129 | 18S952R_MiSeq_0130 | 18S952R_MiSeq_0131 | 18S952R_MiSeq_0132 | 18S952R_MiSeq_0133 | 18S952R_MiSeq_0134 | 18S952R_MiSeq_0135 | 18S952R_MiSeq_0136 | |
| C | RT-03 | PM-02 | BA-03 | CB-03 | RC-03 | HS-03 | OK-03 | NO-03 | NA-03 | VN-03 | PL-03 | AW-03 | |
| | 18S952R_MiSeq_0137 | 18S952R_MiSeq_0138 | 18S952R_MiSeq_0139 | 18S952R_MiSeq_0140 | 18S952R_MiSeq_0141 | 18S952R_MiSeq_0142 | 18S952R_MiSeq_0143 | 18S952R_MiSeq_0144 | 18S952R_MiSeq_0145 | 18S952R_MiSeq_0146 | 18S952R_MiSeq_0147 | 18S952R_MiSeq_0148 | |
| D | RT-04 | PM-03 | BA-04 | CB-04 | RC-04 | HS-04 | OK-04 | NO-04 | NA-04 | VN-04 | PL-04 | AW-04 | |
| | 18S952R_MiSeq_0149 | 18S952R_MiSeq_0150 | 18S952R_MiSeq_0151 | 18S952R_MiSeq_0152 | 18S952R_MiSeq_0153 | 18S952R_MiSeq_0154 | 18S952R_MiSeq_0155 | 18S952R_MiSeq_0156 | 18S952R_MiSeq_0157 | 18S952R_MiSeq_0158 | 18S952R_MiSeq_0159 | 18S952R_MiSeq_0160 | |
| E | RT-05 | PM-04 | BA-05 | CB-05 | RC-05 | HS-05 | OK-05 | NO-05 | NA-05 | VN-05 | PL-05 | AW-05 | |
| | 18S952R_MiSeq_0161 | 18S952R_MiSeq_0162 | 18S952R_MiSeq_0163 | 18S952R_MiSeq_0164 | 18S952R_MiSeq_0165 | 18S952R_MiSeq_0166 | 18S952R_MiSeq_0167 | 18S952R_MiSeq_0168 | 18S952R_MiSeq_0169 | 18S952R_MiSeq_0170 | 18S952R_MiSeq_0171 | 18S952R_MiSeq_0172 | |
| F | | PM-DW | BA-DW | CB-DW | RC-DW | HS-DW | OK-DW | NO-DW | NA-DW | VN-DW | PL-DW | | |
| | 18S952R_MiSeq_0173 | 18S952R_MiSeq_0174 | 18S952R_MiSeq_0175 | 18S952R_MiSeq_0176 | 18S952R_MiSeq_0177 | 18S952R_MiSeq_0178 | 18S952R_MiSeq_0179 | 18S952R_MiSeq_0180 | 18S952R_MiSeq_0181 | 18S952R_MiSeq_0182 | 18S952R_MiSeq_0183 | 18S952R_MiSeq_0184 | |
| G | RT-XB | PM-LM | BA-LM | CB-LM | RC-LM | HS-LM | OK-LM | NO-LM | NA-LM | VN-LM | PL-LM | AW-XB | |
| | 18S952R_MiSeq_0185 | 18S952R_MiSeq_0186 | 18S952R_MiSeq_0187 | 18S952R_MiSeq_0188 | 18S952R_MiSeq_0189 | 18S952R_MiSeq_0190 | 18S952R_MiSeq_0191 | 18S952R_MiSeq_0192 | 18S952R_MiSeq_0193 | 18S952R_MiSeq_0194 | 18S952R_MiSeq_0195 | 18S952R_MiSeq_0196 | |
| H | RT-NG | PM-NG | BA-NG | CB-NG | RC-NG | HS-NG | OK-NG | NO-NG | NA-NG | fw_mock | sw_mock | zebra | |
| | 18S952R_MiSeq_0197 | 18S952R_MiSeq_0198 | 18S952R_MiSeq_0199 | 18S952R_MiSeq_0200 | 18S952R_MiSeq_0201 | 18S952R_MiSeq_0202 | 18S952R_MiSeq_0203 | 18S952R_MiSeq_0204 | 18S952R_MiSeq_0205 | 18S952R_MiSeq_0037 | 18S952R_MiSeq_0038 | 18S952R_MiSeq_0039 | |

c. Cornell Run 2

| | Rotterdam | Puerto Madryn | Buenos Aires | Coos Bay | Richmond | Haines | Oakland | New Orleans | Nanaimo | Vancouver | Portland | Antwerp | |
|---|---|---|---|---|---|---|---|---|---|---|---|---|---|
| | 1 | 2 | 3 | 4 | 5 | 6 | 7 | 8 | 9 | 10 | 11 | 12 | |
| A | RT-01 | PM-01 | BA-01 | CB-01 | RC-01 | HS-01 | OK-01 | NO-01 | NA-01 | VN-01 | PL-01 | AW-01 | sample |
| | 18S952R_MiSeq_0113 | 18S952R_MiSeq_0114 | 18S952R_MiSeq_0115 | 18S952R_MiSeq_0116 | 18S952R_MiSeq_0117 | 18S952R_MiSeq_0118 | 18S952R_MiSeq_0119 | 18S952R_MiSeq_0120 | 18S952R_MiSeq_0121 | 18S952R_MiSeq_0122 | 18S952R_MiSeq_0123 | 18S952R_MiSeq_0124 | barcode |
| B | RT-02 | PM-02 | BA-02 | CB-02 | RC-02 | HS-02 | OK-02 | NO-02 | NA-02 | VN-02 | PL-02 | AW-02 | |
| | 18S952R_MiSeq_0125 | 18S952R_MiSeq_0126 | 18S952R_MiSeq_0127 | 18S952R_MiSeq_0128 | 18S952R_MiSeq_0129 | 18S952R_MiSeq_0130 | 18S952R_MiSeq_0131 | 18S952R_MiSeq_0132 | 18S952R_MiSeq_0133 | 18S952R_MiSeq_0134 | 18S952R_MiSeq_0135 | 18S952R_MiSeq_0136 | |
| C | RT-03 | PM-02 | BA-03 | CB-03 | RC-03 | HS-03 | OK-03 | NO-03 | NA-03 | VN-03 | PL-03 | AW-03 | |
| | 18S952R_MiSeq_0137 | 18S952R_MiSeq_0138 | 18S952R_MiSeq_0139 | 18S952R_MiSeq_0140 | 18S952R_MiSeq_0141 | 18S952R_MiSeq_0142 | 18S952R_MiSeq_0143 | 18S952R_MiSeq_0144 | 18S952R_MiSeq_0145 | 18S952R_MiSeq_0146 | 18S952R_MiSeq_0147 | 18S952R_MiSeq_0148 | |
| D | RT-04 | PM-03 | BA-04 | CB-04 | RC-04 | HS-04 | OK-04 | NO-04 | NA-04 | VN-04 | PL-04 | AW-04 | |
| | 18S952R_MiSeq_0149 | 18S952R_MiSeq_0150 | 18S952R_MiSeq_0151 | 18S952R_MiSeq_0152 | 18S952R_MiSeq_0153 | 18S952R_MiSeq_0154 | 18S952R_MiSeq_0155 | 18S952R_MiSeq_0156 | 18S952R_MiSeq_0157 | 18S952R_MiSeq_0158 | 18S952R_MiSeq_0159 | 18S952R_MiSeq_0160 | |
| E | RT-05 | PM-04 | BA-05 | CB-05 | RC-05 | HS-05 | OK-05 | NO-05 | NA-05 | VN-05 | PL-05 | AW-05 | |
| | 18S952R_MiSeq_0161 | 18S952R_MiSeq_0162 | 18S952R_MiSeq_0163 | 18S952R_MiSeq_0164 | 18S952R_MiSeq_0165 | 18S952R_MiSeq_0166 | 18S952R_MiSeq_0167 | 18S952R_MiSeq_0168 | 18S952R_MiSeq_0169 | 18S952R_MiSeq_0170 | 18S952R_MiSeq_0171 | 18S952R_MiSeq_0172 | |
| F | | PM-DW | BA-DW | CB-DW | RC-DW | HS-DW | OK-DW | NO-DW | NA-DW | VN-DW | PL-DW | | |
| | 18S952R_MiSeq_0173 | 18S952R_MiSeq_0174 | 18S952R_MiSeq_0175 | 18S952R_MiSeq_0176 | 18S952R_MiSeq_0177 | 18S952R_MiSeq_0178 | 18S952R_MiSeq_0179 | 18S952R_MiSeq_0180 | 18S952R_MiSeq_0181 | 18S952R_MiSeq_0182 | 18S952R_MiSeq_0183 | 18S952R_MiSeq_0184 | |
| G | RT-XB | PM-LM | BA-LM | CB-LM | RC-LM | HS-LM | OK-LM | NO-LM | NA-LM | VN-LM | PL-LM | AW-XB | |
| | 18S952R_MiSeq_0185 | 18S952R_MiSeq_0186 | 18S952R_MiSeq_0187 | 18S952R_MiSeq_0188 | 18S952R_MiSeq_0189 | 18S952R_MiSeq_0190 | 18S952R_MiSeq_0191 | 18S952R_MiSeq_0192 | 18S952R_MiSeq_0193 | 18S952R_MiSeq_0194 | 18S952R_MiSeq_0195 | 18S952R_MiSeq_0196 | |
| H | RT-NG | PM-NG | BA-NG | CB-NG | RC-NG | HS-NG | OK-NG | NO-NG | NA-NG | fw_mock | sw_mock | zebra | |
| | 18S952R_MiSeq_0197 | 18S952R_MiSeq_0198 | 18S952R_MiSeq_0199 | 18S952R_MiSeq_0200 | 18S952R_MiSeq_0201 | 18S952R_MiSeq_0202 | 18S952R_MiSeq_0203 | 18S952R_MiSeq_0204 | 18S952R_MiSeq_0205 | 18S952R_MiSeq_0037 | 18S952R_MiSeq_0038 | 18S952R_MiSeq_0039 | |

d. Cornell Run 3

| extraction: details: | columns 1 - 4 | | | | Milne Inlet 31.07.2018 book 2 page 2 | | | | | Wilmington 16.07.2018 book 2 page 8 | Ghent 04.12.2017 book 2 page 8 | Zeebrugge 04.12.2017 book 2 page 8 | |
|---|---|---|---|---|---|---|---|---|---|---|---|---|---|
| | 1 | 2 | 3 | 4 | 5 | 6 | 7 | 8 | 9 | 10 | 11 | 12 | sample |
| A | | | | | ML-01 | ML-08 | ML-15 | ML-22 | ML-29 | WL-01 | GH-01 | ZB-01 | barcode |
| | 18S952R_MiSeq_0209 | 18S952R_MiSeq_0210 | 18S952R_MiSeq_0211 | 18S952R_MiSeq_0212 | 18S952R_MiSeq_0213 | 18S952R_MiSeq_0214 | 18S952R_MiSeq_0215 | 18S952R_MiSeq_0216 | 18S952R_MiSeq_0217 | 18S952R_MiSeq_0218 | 18S952R_MiSeq_0219 | 18S952R_MiSeq_0220 | |
| B | | | | | ML-02 | ML-09 | ML-16 | ML-23 | ML-30 | WL-02 | GH-02 | ZB-02 | |
| | 18S952R_MiSeq_0221 | 18S952R_MiSeq_0222 | 18S952R_MiSeq_0223 | 18S952R_MiSeq_0224 | 18S952R_MiSeq_0225 | 18S952R_MiSeq_0226 | 18S952R_MiSeq_0227 | 18S952R_MiSeq_0228 | 18S952R_MiSeq_0229 | 18S952R_MiSeq_0230 | 18S952R_MiSeq_0231 | 18S952R_MiSeq_0232 | |
| C | | | | | ML-03 | ML-10 | ML-17 | ML-24 | ML-31 | WL-03 | GH-03 | ZB-03 | |
| | 18S952R_MiSeq_0233 | 18S952R_MiSeq_0234 | 18S952R_MiSeq_0235 | 18S952R_MiSeq_0236 | 18S952R_MiSeq_0237 | 18S952R_MiSeq_0238 | 18S952R_MiSeq_0239 | 18S952R_MiSeq_0240 | 18S952R_MiSeq_0241 | 18S952R_MiSeq_0242 | 18S952R_MiSeq_0243 | 18S952R_MiSeq_0244 | |
| D | | | | | ML-04 | ML-11 | ML-18 | ML-25 | ML-DW1 | WL-04 | GH-04 | ZB-04 | |
| | 18S952R_MiSeq_0245 | 18S952R_MiSeq_0246 | 18S952R_MiSeq_0247 | 18S952R_MiSeq_0248 | 18S952R_MiSeq_0249 | 18S952R_MiSeq_0250 | 18S952R_MiSeq_0251 | 18S952R_MiSeq_0252 | 18S952R_MiSeq_0253 | 18S952R_MiSeq_0254 | 18S952R_MiSeq_0255 | 18S952R_MiSeq_0256 | |
| E | | | | | ML-05 | ML-12 | ML-19 | ML-26 | ML-DW2 | WL-05 | GH-05 | ZB-05 | |
| | 18S952R_MiSeq_0257 | 18S952R_MiSeq_0258 | 18S952R_MiSeq_0259 | 18S952R_MiSeq_0260 | 18S952R_MiSeq_0261 | 18S952R_MiSeq_0262 | 18S952R_MiSeq_0263 | 18S952R_MiSeq_0264 | 18S952R_MiSeq_0265 | 18S952R_MiSeq_0266 | 18S952R_MiSeq_0267 | 18S952R_MiSeq_0268 | |
| F | | | | | ML-06 | ML-13 | ML-20 | ML-27 | ML-NG | WL-DW | GH-NG | ZB-NG | |
| | 18S952R_MiSeq_0269 | 18S952R_MiSeq_0270 | 18S952R_MiSeq_0271 | 18S952R_MiSeq_0272 | 18S952R_MiSeq_0273 | 18S952R_MiSeq_0274 | 18S952R_MiSeq_0275 | 18S952R_MiSeq_0276 | 18S952R_MiSeq_0277 | 18S952R_MiSeq_0278 | 18S952R_MiSeq_0279 | 18S952R_MiSeq_0280 | |
| G | | | | | ML-07 | ML-14 | ML-21 | ML-28 | ML-NG | WL-XB | GH-XB | ZB-XB | |
| | 18S952R_MiSeq_0281 | 18S952R_MiSeq_0282 | 18S952R_MiSeq_0283 | 18S952R_MiSeq_0284 | 18S952R_MiSeq_0285 | 18S952R_MiSeq_0286 | 18S952R_MiSeq_0287 | 18S952R_MiSeq_0288 | 18S952R_MiSeq_0289 | 18S952R_MiSeq_0290 | 18S952R_MiSeq_0291 | 18S952R_MiSeq_0292 | |
| H | | | | | | | | | | fw_mock | sw_mock | zebra | |
| | 18S952R_MiSeq_0293 | 18S952R_MiSeq_0294 | 18S952R_MiSeq_0295 | 18S952R_MiSeq_0296 | 18S952R_MiSeq_0297 | 18S952R_MiSeq_0298 | 18S952R_MiSeq_0299 | 18S952R_MiSeq_0300 | 18S952R_MiSeq_0301 | 18S952R_MiSeq_0037 | 18S952R_MiSeq_0038 | 18S952R_MiSeq_0039 | |

**Figure S6**

Read counts at each stage of the bioinformatics workflow [purple = input raw read counts, turquoise = read counts after quality filtering, blue = recounts after denoising, and yellow = read counts after merging paired ends] for samples from (a) Singapore, (b) Adelaide, (c) Chicago, and (d) the remaining ports. Singapore, Chicago and Adelaide samples were sequenced separately for another study (Grey et al. 2018). Ports are denoted in figure (d) as two-letter abbreviations at the beginning of each sample name as follows: Antwerp (AW), Baltimore (BA), Coos Bay (CB), Ghent (GH), Haines (HS), Honolulu (HN), Houston (HT), Long Beach (LB), Miami (MI), New Orleans (NO), Oakland (OK), Portland (PL), Puerto Madryn (PM), Richmond (RC), Rotterdam (RT), Wilmington (WL), and Zeebrugge (ZB).

a. Singapore

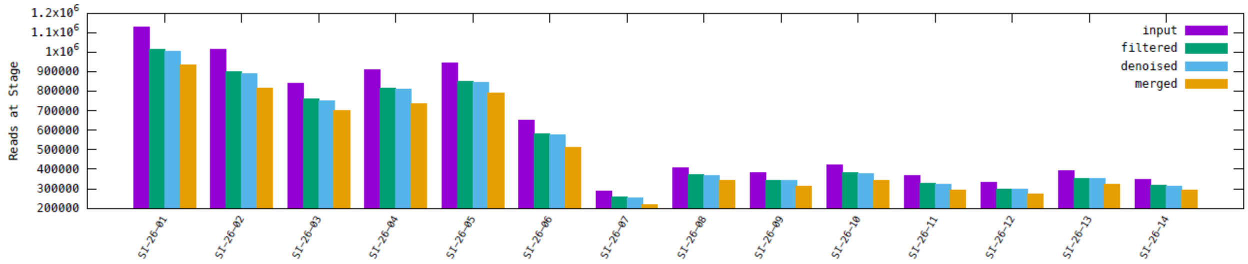


b. Adelaide

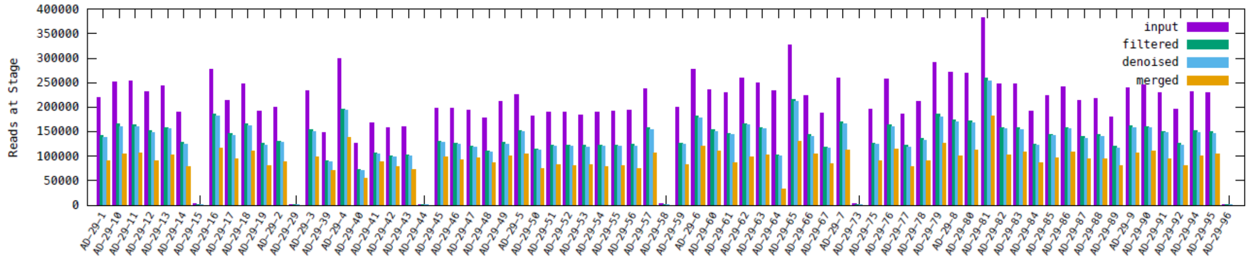


c. Chicago

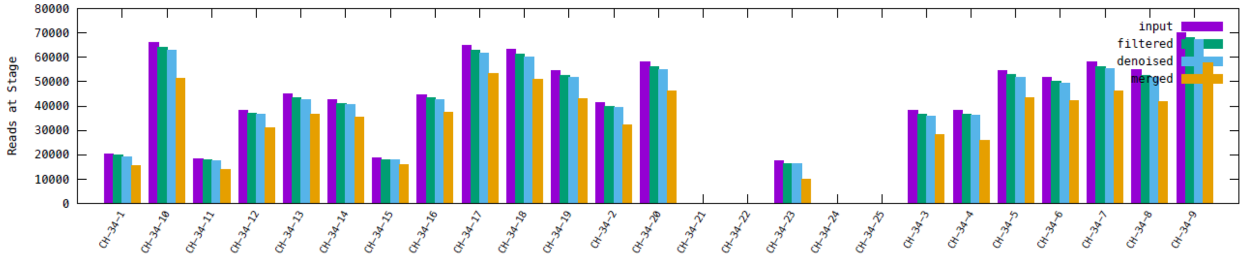


d. Remaining ports (see legend for port abbreviations)

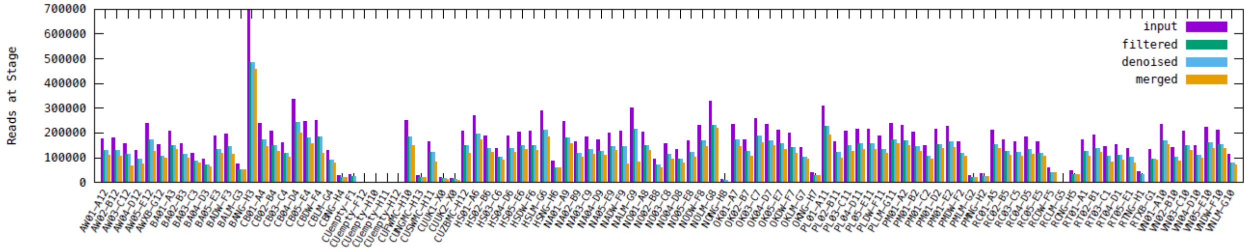

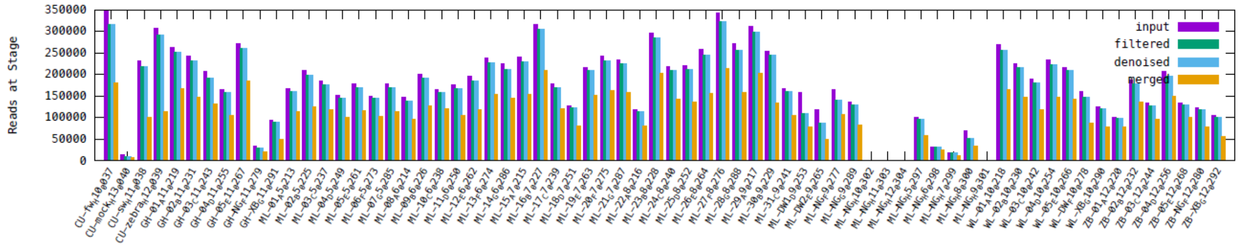
Reads at Stage
350000
300000
250000
200000
150000
100000
50000
0
input
filtered
denoised
merged

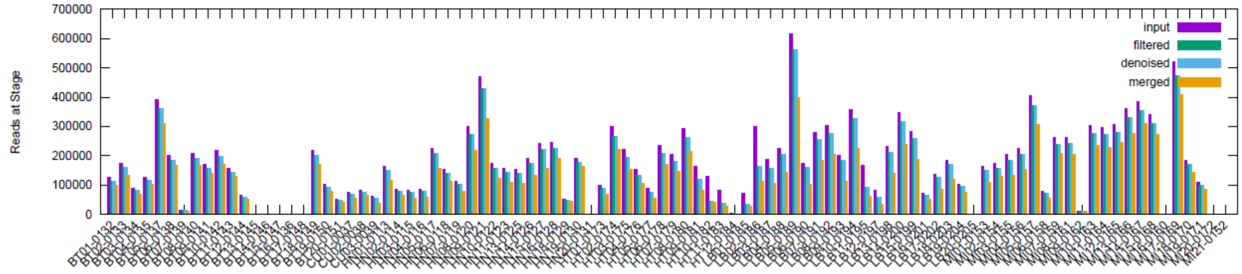
Reads at Stage
700000
600000
500000
400000
300000
200000
100000
0
input
filtered
denoised
merged

**Fig. S7.**
Accumulation of unique Amplicon Sequence Variants (ASVs) among increasing amounts of random sequences drawn from each port. Shown are mean values and 95% confidence intervals between three replicates. Dotted line indicates a rarefaction depth of 37900, dashed line a rarefaction depth of 49900. Port abbreviations: AD – Adelaide, AW – Antwerp, BT – Baltimore, CB – Coos-Bay, GH – Ghent, HN – Honolulu, HN – Haines, HT – Houston, LB – Long Beach, MI – Miami, NA – Nanaimo, NO – New-Orleans, OK – Oakland, PL – Portland, PM – Puerto-Madryn, RT – Richmond, RT – Rotterdam, SI – Singapore, VN – Vancouver, WL – Wilmington, ZB – Zeebrugge.

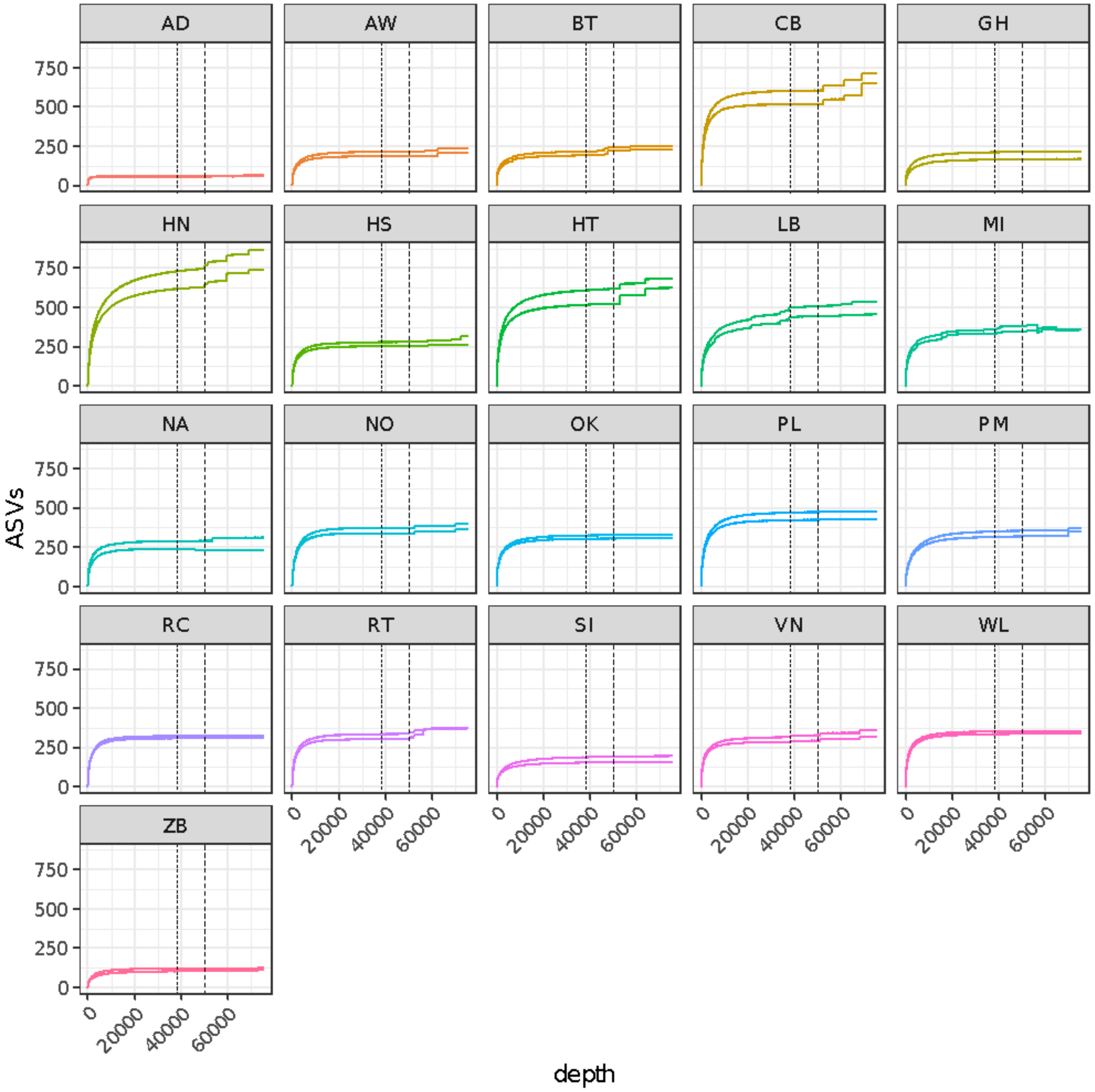

**Fig. S8.**

Unifrac distances decay among 9 port-pairs as a function of sampling effort (1-7 samples from each port in the pair) from the deeply rarefied dataset. These 9 port pairs were randomly selected to infer the number of samples needed to generate robust Unifrac estimates. Port abbreviations: AD – Adelaide, AW – Antwerp, BT – Baltimore, CB – Coos-Bay, GH – Ghent, HN – Honolulu, HN – Haines, HT – Houston, LB – Long Beach, MI – Miami, NA – Nanaimo, NO – New-Orleans, OK – Oakland, PL – Portland, PM – Puerto-Madryn, RT – Richmond, RT – Rotterdam, SI – Singapore, VN – Vancouver, WL – Wilmington, ZB – Zeebrugge.

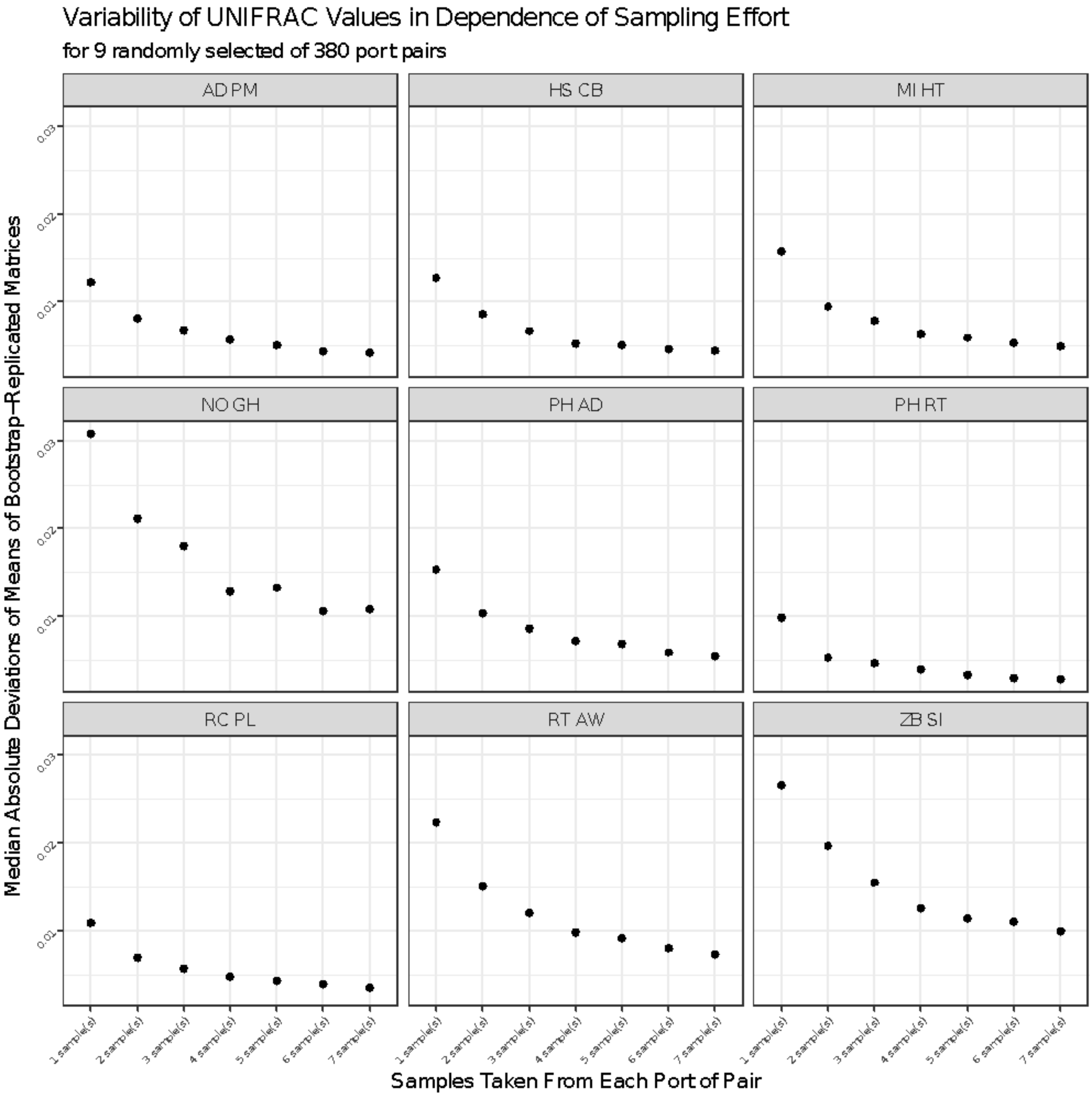

**Table S1. Port sample dates, times, and coordinates.**

| Port | Date | Local Time | Latitude | Longitude |
|---|---|---|---|---|
| Puerto Madryn, ARG | 2018-10-10 | 8:00 | -42.7445 | -65.0367 |
| Adelaide, AUS | 2014-07-03 | 11:00 | -34.8172 | 138.5100 |
| Singapore, SGP | 2014-07-24 | 13:30 | 1.4549 | 103.7783 |
| Honolulu, USA | 2016-11-22 | 10:20 | 21.3039 | -157.8706 |
| Miami, USA | 2016-06-16 | 11:51 | 25.7677 | -80.1656 |
| Houston, USA | 2016-06-15 | 11:57 | 29.7618 | -95.0821 |
| New Orleans, USA | 2016-06-09 | 13:18 | 29.9218 | -90.1352 |
| Long Beach, USA | 2016-10-12 | 13:50 | 33.7431 | -118.2075 |
| Oakland, CA USA | 2016-07-11 | 15:06 | 37.7933 | -122.2969 |
| Richmond, CA, USA | 2016-07-11 | 9:48 | 37.9064 | -122.3636 |
| Baltimore, USA | 2016-06-09 | 10:26 | 39.2613 | -76.6007 |
| Wilmington, DE USA | 2016-06-27 | 11:13 | 39.7210 | -75.5294 |
| Chicago, USA* | 2013-11-20 | 13:00 | 41.6648 | -87.5879 |
| Coos Bay, USA | 2016-09-27 | 12:22 | 43.3466 | -124.3207 |
| Portland, OR USA | 2016-08-25 | 8:36 | 45.6411 | -122.7847 |
| Nanaimo, CAN* | 2016-09-26 | 10:30 | 49.1713 | -123.9348 |

| | | | | |
|---|---|---|---|---|
| Vancouver, CAN* | 2016-10-22 | 11:14 | 49.2748 | -123.1853 |
| Ghent, BEL | 2018-06-25 | 9:53 | 51.0848 | 3.7340 |
| Antwerp, BEL | 2018-06-27 | 16:00 | 51.2731 | 4.2482 |
| Zeebrugge, BEL | 2018-06-26 | 13:40 | 51.3184 | 3.2194 |
| Rotterdam, NLD | 2018-07-10 | 12:00 | 51.8917 | 4.3982 |
| Haines, USA | 2016-08-16 | 3:30 | 59.2278 | -135.4364 |

**Samples from these ports are not included in deep rarefaction analyses.*

**Table S2. Coefficients of the AIC-selected models (deep rarefaction of samples from 19 ports).**

<u>Direct traffic risk:</u> <u>UNIFRAC = ENV + BGR + ENV*BGR+ (ORG) + (DEST)</u>

Fixed effects:

| | Estimate | Std. Error |
|---|---|---|
| (Intercept) | 0.758549 | 0.014309 |
| ENV | 0.061923 | 0.011743 |
| BRG | 0.008963 | 0.011344 |
| ENV*BGR | -0.034278 | 0.012600 |

<u>Stepping-stone traffic risk:</u> <u>UNIFRAC = ENV+ SHP + ENV*SHP + (ORG) + (DEST)</u>

Fixed effects:

| | Estimate | Std. Error |
|---|---|---|
| (Intercept) | 0.766606 | 0.011204 |
| ENV | 0.030477 | 0.005341 |
| SHP | -0.002653 | 0.004804 |
| ENV*BGR | 0.013167 | 0.004396 |

<u>Direct ballast risk:</u> <u>UNIFRAC = ENV + BGR+ SHP + BGR*ENV+BGR*SHP + (ORG) + (DEST)</u>

Fixed effects:

| | Estimate | Std. Error |
|---|---|---|
| (Intercept) | 0.7693570 | 0.0174270 |
| ENV | 0.0620989 | 0.0167782 |
| BRG | 0.0004407 | 0.0137807 |
| SHP | -0.0085444 | 0.0076253 |
| ENV*BRG | -0.0342781 | 0.0156476 |
| ENV*SHP | -0.0005381 | 0.0084644 |

<u>Stepping-stone ballast risk:</u> <u>UNIFRAC = ENV + SHP + ENV*SHP+ (ORG) + (DEST)</u>

Fixed effects:

| | Estimate | Std. Error |
|---|---|---|
| (Intercept) | 0.766456 | 0.011207 |
| ENV | 0.030581 | 0.005345 |
| SHP | -0.002999 | 0.004790 |
| ENV*SHP | 0.012538 | 0.004403 |

**Table S3.** AIC-Selected models for both deep- and shallow-rarefied datasets. Orange row indicates the model with the lowest AIC.

| | DEEP RAREFACTION (n= 19 PORTS) | | | | | |
|---|---|---|---|---|---|---|
| **Model** | **Model family** | **Connection Type** | **Traffic-related Variable** | **SELECTED MODEL** | **AIC** | **BIC** |
| model.FreqTrip | Transport frequency | Direct | VOY_FREQ | RESP_UNIFRAC ~ PRED_ENV + ECO_DIFF : PRED_ENV + ECO_DIFF + (1 \| PORT) + (1 \| DEST) | -476.9 | -454.91 |
| model.JFreqTrip | Transport frequency | Stepping stone | J_VOY_FREQ | RESP_UNIFRAC ~ PRED_EN + **J_VOY_FREQ** + PRED_ENV : J_VOY_FREQ + (1\|PORT) + (1\|DEST) | -479.34 | -457.35 |
| model.B_F | Transport Risk | Direct | B_**FON**_NOECO_NOENV | RESP_UNIFRAC ~ PRED_ENV + ECO_DIFF + B_FON_NOECO_NOENV + ECO_DIFF:PRED_ENV + ECO_DIFF : B_FON_NOECO_NOEN + (1\|PORT) + (1\|DEST) | -478.51 | -451.23 |
| model.JAC_B_F | Transport Risk | Stepping stone | J_B_**FON**_NOECO_NOENV | RESP_UNIFRAC ~ PRED_ENV + J_B_**FON**_NOECO_NOENV + PRED_EN:J_B_FON_NOECO_NOENV+ (1 \| PORT) + (1 \| DEST) | -478.78 | -456.78 |

| | SHALLOW RAREFACTION (n= 22 PORTS) | | | | | |
|---|---|---|---|---|---|---|
| **Model** | **Model family** | **Connection Type** | **Traffic-related Variable** | **SELECTED MODEL** | **AIC** | **BIC** |
| model.FreqTrip | Transport frequency | Direct | VOY_FREQ | RESP_UNIFRAC ~ PRED_ENV + ECO_DIFF : PRED_ENV + ECO_DIFF + (1 \| PORT) + (1 \| DEST) | -574.55 | -541.12 |
| model.JFreqTrip | Transport frequency | Stepping stone | J_VOY_FREQ | RESP_UNIFRAC ~ ECO_DIFF + PRED_ENV + **J_VOY_FREQ** + ECO_DIFF : J_VOY_FREQ + PRED_ENV : J_VOY_FREQ +(1\|PORT) + (1\|DEST) | -576.42 | -546.3 |
| model.JAC_B_F | Transport Risk | Direct | J_B_**FON**_NOECO_NOENV | RESP_UNIFRAC ~ ECO_DIFF + PRED_ENV **+J_B_FON_NOECO_NOENV** + ECO_DIFF : J_B_FON_NOECO_NOENV + PRED_ENV : J_B_FON_NOECO_NOENV + (1\|PORT) + (1\|DEST) | -575.67 | -545.54 |
| model.B_F | Transport Risk | Stepping stone | B_**FON**_NOECO_NOENV | RESP_UNIFRAC~ ECO_DIFF + PRED_ENV **+B_FON_NOECO_NOENV** + | -571.14 | -541.02 |

**Table S4.**

Non-indigenous species read counts by port. Includes only ASVs unambiguously matched to known non-indigenous species as described in the main text. Species ranges from sources noted in the last column. Observations that represent known native ranges are shaded light green, native range extensions in dark green, known non-native ranges in orange, and new non-native occurrences in red. Port abbreviations: AD – Adelaide, AW – Antwerp, BT – Baltimore, CB – Coos-Bay, GH – Ghent, HN – Honolulu, HN – Haines, HT – Houston, LB – Long Beach, MI – Miami, NA – Nanaimo, NO – New-Orleans, OK – Oakland, PL – Portland, PM – Puerto-Madryn, RT – Richmond, RT – Rotterdam, SI – Singapore, WL – Wilmington, ZB – Zeebrugge.

| Class | Species | ASVs | PH | SI | AD | BT | HN | HT | LB | MI | AW | CB | HS | NO | OK | PL | PM | RC | RT | GH | WL | ZB | Native Range | Non-Native Range |
|---|---|---|---|---|---|---|---|---|---|---|---|---|---|---|---|---|---|---|---|---|---|---|---|---|
| Anthozoa | Diadumene leucolena | 1 | 0 | 0 | 0 | 4926 | 0 | 0 | 0 | 0 | 0 | 0 | 0 | 0 | 0 | 0 | 0 | 0 | 0 | 0 | 0 | 0 | northwest Atlantic | Caribbean, northeast Atlantic, Hawaii, Tropical east Atlantic |
| Anthozoa | Diadumene lineata | 1 | 0 | 0 | 0 | 0 | 0 | 0 | 0 | 0 | 0 | 0 | 0 | 0 | 0 | 0 | 110 | 0 | 0 | 0 | 0 | 0 | northwest Pacific | northeast Pacific, Atlantic, Mediterranean, Black Sea, IndoPacific, New Zealand, Argentina |
| Anthozoa | Exaiptasia pallida | 1 | 0 | 0 | 0 | 0 | 29 | 0 | 0 | 0 | 0 | 0 | 0 | 0 | 0 | 0 | 0 | 0 | 0 | 0 | 0 | 0 | northwest Atlantic, Pacific | northeast Atlantic |
| Ascidia | Ascidia ceratodes | 3 | 103 | 0 | 0 | 0 | 0 | 0 | 5445 | 0 | 0 | 0 | 0 | 0 | 0 | 0 | 0 | 0 | 0 | 0 | 0 | 0 | east Pacific | |
| Ascidia | Ascidia sydneiensis samea | 1 | 108 | 0 | 0 | 0 | 240 | 0 | 0 | 0 | 0 | 0 | 0 | 0 | 0 | 0 | 0 | 0 | 0 | 0 | 0 | 0 | IndoPacific | Hawaii/Guam, Caribbean, Florida |
| Ascidia | Botrylloides leachii | 1 | 0 | 0 | 0 | 0 | 0 | 0 | 0 | 0 | 0 | 0 | 0 | 0 | 14 | 0 | 0 | 68 | 0 | 0 | 0 | 0 | northeast Atlantic | New Zealand, posssibly Australia |
| Ascidia | Ciona intestinalis | 8 | 0 | 0 | 0 | 0 | 0 | 0 | 0 | 0 | 0 | 0 | 0 | 0 | 987 | 0 | 0 | 4133 | 0 | 0 | 0 | 0 | north Atlantic | south Atlantic, Pacific, Indian Ocean |
| Ascidia | Ciona savignyi | 41 | 0 | 0 | 0 | 0 | 0 | 0 | 0 | 0 | 0 | 0 | 0 | 0 | 0 | 0 | 0 | 2721 | 0 | 0 | 0 | 0 | northwest Pacific | northeast Pacific, southwest Atlantic |
| Ascidia | Corella eumyota | 1 | 0 | 0 | 0 | 0 | 0 | 0 | 0 | 0 | 0 | 0 | 0 | 0 | 0 | 0 | 327 | 0 | 0 | 0 | 0 | 0 | subAntarctic, cosmopolitan | northeast Atlantic, likely southern stowaway to southern hemisphere |
| Ascidia | Microcosmus exasperatus | 1 | 13 | 0 | 0 | 0 | 75 | 0 | 0 | 0 | 0 | 0 | 0 | 0 | 0 | 0 | 0 | 0 | 0 | 0 | 0 | 0 | cosmopolitan | Meditteranean, Red Sea |
| Ascidia | Molgula manhattensis | 1 | 0 | 0 | 0 | 0 | 0 | 0 | 0 | 0 | 0 | 0 | 0 | 0 | 199 | 0 | 0 | 35 | 0 | 0 | 0 | 0 | northwest Atlantic | northeast Atlantic, Japan, Australia, Argentina, northeast Pacific |
| Ascidia | Styela canopus | 2 | 0 | 644 | 0 | 0 | 160 | 0 | 0 | 0 | 0 | 0 | 0 | 0 | 0 | 0 | 0 | 0 | 0 | 0 | 0 | 0 | IndoPacific | east Pacific (Mazatlan - Los Angeles), Hawaii & Guam, west Atlantic (Brazil - Nova Scotia), east Atlantic (Senegal - England), Egypt, Lebanon, Adriatic Sea |
| Ascidia | Styela clava | 1 | 0 | 0 | 0 | 0 | 0 | 0 | 98 | 0 | 0 | 0 | 0 | 0 | 124 | 0 | 0 | 62 | 0 | 0 | 0 | 48 | northwest Pacific | northeast Pacific, north Atlantic, Black Sea, Australia, New Zealand, Argentina |
| Ascidia | Styela plicata | 1 | 0 | 0 | 0 | 0 | 28 | 0 | 0 | 0 | 0 | 0 | 0 | 0 | 0 | 0 | 0 | 0 | 0 | 0 | 0 | 0 | cosmopolitan, northwest Pacific | Gulf of Mexico, California, Australia, New Zealand, southwest Atlantic |
| Bacillariophyceae | Chaetoceros didymus | 1 | 0 | 0 | 0 | 0 | 0 | 0 | 0 | 0 | 0 | 0 | 0 | 0 | 0 | 0 | 0 | 2 | 0 | 0 | 0 | 0 | cosmopolitan | Mexico |
| Bacillariophyceae | Chaetoceros peruvianus | 1 | 557 | 0 | 0 | 0 | 0 | 0 | 16 | 0 | 0 | 0 | 0 | 0 | 0 | 0 | 0 | 0 | 0 | 0 | 0 | 0 | cosmopolitanl | Black Sea |
| Bacillariophyceae | Conticribra guillardii | 2 | 0 | 0 | 0 | 18 | 0 | 0 | 0 | 0 | 17 | 0 | 1930 | 117 | 53 | 0 | 0 | 128 | 135 | 0 | 344 | 0 | cosmopolitan | Great Lakes, Poland |
| Bacillariophyceae | Conticribra weissflogii | 8 | 0 | 0 | 0 | 0 | 0 | 108 | 0 | 0 | 0 | 0 | 0 | 0 | 0 | 48675 | 0 | 0 | 75 | 0 | 0 | 0 | cosmopolitan; possibly Baltic | |
| Bacillariophyceae | Ditylum brightwellii | 1 | 0 | 0 | 0 | 0 | 0 | 0 | 0 | 0 | 0 | 221 | 0 | 0 | 0 | 0 | 0 | 0 | 0 | 0 | 0 | 168 | cosmopolitan | |
| Bacillariophyceae | Pleurosira laevis | 2 | 0 | 0 | 0 | 0 | 0 | 37 | 0 | 0 | 0 | 0 | 0 | 0 | 0 | 0 | 0 | 0 | 0 | 0 | 0 | 0 | cosmopolitan | Great Lakes, Europe |
| Bacillariophyceae | Pseudo-nitzschia delicatissima | 1 | 0 | 0 | 0 | 0 | 0 | 0 | 294 | 0 | 0 | 0 | 0 | 0 | 0 | 0 | 0 | 4066 | 0 | 0 | 0 | 0 | cosmopolitan | Mexico |
| Bacillariophyceae | Pseudo-nitzschia multistriata | 2 | 193 | 0 | 0 | 0 | 83 | 0 | 332 | 0 | 0 | 0 | 0 | 0 | 388 | 0 | 0 | 219 | 0 | 0 | 0 | 0 | cosmopolitan | Italy |
| Bacillariophyceae | Pseudo-nitzschia pseudodelicatissima | 1 | 0 | 0 | 0 | 0 | 0 | 0 | 0 | 0 | 0 | 6 | 0 | 0 | 0 | 0 | 0 | 0 | 0 | 0 | 0 | 0 | cosmopolitan | Mexico |
| Bacillariophyceae | Pseudo-nitzschia pungens | 1 | 29 | 0 | 0 | 0 | 0 | 0 | 0 | 0 | 0 | 426 | 0 | 0 | 0 | 0 | 0 | 0 | 0 | 0 | 0 | 0 | cosmopolitan | Mexico |
| Bacillariophyceae | Rhizosolenia setigera | 1 | 99 | 0 | 0 | 0 | 0 | 0 | 0 | 0 | 0 | 0 | 0 | 0 | 0 | 0 | 0 | 0 | 0 | 0 | 0 | 0 | cosmopolitan | Ukraine |
| Bacillariophyceae | Skeletonema potamos | 3 | 0 | 0 | 0 | 383 | 0 | 13024 | 0 | 0 | 0 | 0 | 0 | 19894 | 0 | 8583 | 0 | 0 | 27465 | 5431 | 75916 | 0 | cosmopolitan | Great Lakes, Czech Republic, Poland, France |
| Bacillariophyceae | Skeletonema subsalsum | 1 | 0 | 0 | 0 | 0 | 0 | 0 | 0 | 0 | 0 | 0 | 0 | 0 | 0 | 0 | 0 | 0 | 111 | 0 | 0 | 0 | northeast Atlantic, Caspian Sea, Black Sea | Great Lakes, Argentina, Brazil, Gulf of Mexico, Ireland |
| Bacillariophyceae | Thalassiosira anguste-lineata | 1 | 0 | 0 | 0 | 0 | 0 | 0 | 0 | 0 | 0 | 0 | 0 | 0 | 0 | 0 | 0 | 0 | 0 | 0 | 0 | 599 | cosmopolitan | Mexico |
| Bivalva | Arcuatula senhousia | 1 | 0 | 1422 | 0 | 0 | 0 | 0 | 0 | 0 | 0 | 0 | 0 | 0 | 7000 | 0 | 0 | 538 | 0 | 0 | 0 | 0 | central & northwest Pacific | southwest Pacific, northeast Pacific, east Atlantic, tropical west Atlantic |
| Bivalva | Brachidontes exustus | 1 | 0 | 0 | 0 | 0 | 0 | 346 | 0 | 0 | 0 | 0 | 0 | 0 | 0 | 0 | 0 | 0 | 0 | 0 | 0 | 0 | Gulf of Mexico, west Atlantic | Ireland/England |
| Bivalva | Clementia papyracea | 1 | 0 | 259 | 0 | 0 | 0 | 0 | 0 | 0 | 0 | 0 | 0 | 0 | 0 | 0 | 0 | 0 | 0 | 0 | 0 | 0 | IndoPacific | Mediterranean |
| Bivalva | Crassostrea virginica | 1 | 0 | 0 | 0 | 0 | 0 | 59 | 0 | 0 | 0 | 0 | 0 | 0 | 0 | 0 | 0 | 0 | 0 | 0 | 0 | 0 | northwest Atlantic | Hawaii |
| Bivalva | Isognomon ephippium | 1 | 0 | 0 | 0 | 0 | 2738 | 0 | 0 | 0 | 0 | 0 | 0 | 0 | 0 | 0 | 0 | 0 | 0 | 0 | 0 | 0 | IndoPacific, Indian Ocean | Mediterreanean |
| Bivalva | Lyrodus pedicellatus | 3 | 0 | 0 | 0 | 0 | 65 | 0 | 0 | 0 | 0 | 0 | 0 | 0 | 0 | 0 | 0 | 0 | 0 | 0 | 0 | 0 | cryptogenic | northeast Pacific, Hawaii, New Zealand, South Africa, southwest Atlantic |
| Bivalva | Mya arenaria | 1 | 0 | 0 | 0 | 0 | 0 | 0 | 0 | 0 | 0 | 0 | 0 | 0 | 0 | 0 | 0 | 0 | 29 | 0 | 0 | 0 | northwest Atlantic, possibly Alaska | northeast Atlantic, Black Sea, northeast Pacific |
| Bivalva | Mytilopsis leucophaeata | 2 | 0 | 0 | 0 | 4680 | 0 | 412 | 0 | 0 | 1623 | 0 | 0 | 0 | 0 | 0 | 0 | 0 | 37 | 8264 | 0 | 16 | northwest Atlantic | New England, northeast Atlantic, Black Sea |
| Bivalva | Mytilus edulis | 2 | 0 | 0 | 937 | 0 | 0 | 0 | 207 | 0 | 0 | 21 | 356 | 0 | 15 | 0 | 1759 | 0 | 0 | 0 | 0 | 366 | north Atlantic, possibly southwest Atlantic c | northeast Pacific, Chile, Argentina, Australia, New Zealand * Note: The Puerto Madrin (Argentina) population of Mytillus is hybrid. Genetics Selection Evolution 50: 5 (2018)). |
| Bivalve | Rangia cuneata | 1 | 0 | 0 | 0 | 0 | 0 | 39 | 0 | 0 | 0 | 0 | 0 | 0 | 0 | 0 | 0 | 0 | 15 | 463 | 0 | 0 | Gulf of Mexico | northwest Atlantic (North Carolina - New York), northeast Atlantic |
| Copepoda | Acartia tonsa | 5 | 0 | 0 | 0 | 22 | 0 | 0 | 0 | 0 | 8712 | 0 | 0 | 0 | 0 | 0 | 0 | 0 | 1139 | 942 | 0 | 0 | cosmopolitan, possible Northeast NA glacial | northeast Atlantic |
| Copepoda | Euterpina acutifrons | 3 | 28 | 0 | 0 | 0 | 0 | 0 | 0 | 0 | 0 | 0 | 0 | 0 | 6 | 0 | 0 | 0 | 0 | 0 | 0 | 0 | cosmopolitan | Columbia |
| Copepoda | Oithona davisae | 6 | 0 | 60017 | 0 | 0 | 0 | 0 | 0 | 0 | 0 | 0 | 0 | 0 | 161 | 0 | 0 | 109 | 0 | 0 | 0 | 104 | northwest Pacific | northeast Pacific, northeast Atlantic, Black Sea, Chile (1 sample) |
| Copepoda | Pseudocalanus elongatus | 9 | 0 | 0 | 0 | 0 | 0 | 0 | 0 | 0 | 0 | 11498 | 0 | 0 | 0 | 0 | 0 | 0 | 0 | 0 | 0 | 0 | north Atlantic, Black Sea | Mediterranean |
| Decapoda | Callinectes sapidus | 1 | 0 | 0 | 0 | 0 | 0 | 29 | 0 | 0 | 0 | 0 | 0 | 0 | 0 | 0 | 0 | 0 | 0 | 0 | 0 | 0 | west Atlantic | east Atlantic, Mediterranean, Adriatic, Black, Japan |
| Decapoda | Carcinus maenas | 1 | 0 | 0 | 0 | 0 | 0 | 0 | 0 | 0 | 0 | 0 | 0 | 0 | 0 | 0 | 20 | 0 | 0 | 0 | 0 | 0 | northeast Atlantic | northwest Atlantic, southwest Atlantic, southeast Atlantic, east Pacific, Australia |
| Demospongiae | Suberites aurantiacus | 1 | 17 | 0 | 0 | 0 | 0 | 0 | 0 | 0 | 0 | 0 | 0 | 0 | 0 | 0 | 0 | 0 | 0 | 0 | 0 | 0 | Caribbean and tropical west Atlantic | Hawaii |
| Hydrozoa | Cordylophora caspia | 3 | 0 | 0 | 0 | 59 | 0 | 398 | 0 | 0 | 0 | 0 | 0 | 0 | 0 | 0 | 0 | 0 | 1892 | 2916 | 1834 | 0 | Black Sea, Caspian Sea | global, multiple lineages |
| Hydrozoa | Coryne eximia | 2 | 0 | 0 | 0 | 0 | 0 | 0 | 0 | 0 | 0 | 7445 | 0 | 0 | 0 | 0 | 0 | 0 | 0 | 0 | 0 | 0 | cosmopolitan | |
| Hydrozoa | Eudendrium carneum | 1 | 0 | 31139 | 0 | 0 | 0 | 0 | 0 | 8 | 0 | 0 | 0 | 0 | 0 | 0 | 0 | 0 | 0 | 0 | 0 | 0 | circumsubtropical | Mediterranean, Indian Ocean |
| Hydrozoa | Gonothyraea loveni | 1 | 0 | 0 | 0 | 0 | 0 | 0 | 0 | 0 | 0 | 0 | 0 | 0 | 7 | 0 | 0 | 0 | 0 | 0 | 0 | 0 | cosmopolitan | South Africa, southwest Atlantic |
| Hydrozoa | Obelia dichotoma | 3 | 4 | 0 | 0 | 0 | 0 | 0 | 0 | 0 | 0 | 0 | 0 | 0 | 0 | 0 | 27 | 0 | 0 | 0 | 0 | 108 | cosmopolitan | South Africa, cryptogenic in southwest Atlantic |
| Hydrozoa | Obelia geniculata | 1 | 0 | 0 | 0 | 0 | 0 | 0 | 0 | 0 | 0 | 24 | 0 | 0 | 0 | 0 | 0 | 0 | 0 | 0 | 0 | 0 | cosmopolitan | South Africa |
| Maxillopoda | Balanus glandula | 1 | 0 | 0 | 0 | 0 | 0 | 0 | 0 | 0 | 0 | 0 | 0 | 0 | 0 | 0 | 25 | 0 | 0 | 0 | 0 | 0 | northeast Pacific | South Africa, Argentina, Japan |
| Polychaeta | Alitta succinea | 1 | 0 | 0 | 0 | 12 | 0 | 0 | 0 | 0 | 0 | 0 | 0 | 0 | 0 | 0 | 0 | 0 | 0 | 0 | 0 | 0 | north Atlantic | Pacific, southwest Atlantic |
| Polychaeta | Ficopomatus enigmaticus | 42 | 0 | 0 | 0 | 0 | 0 | 0 | 0 | 0 | 33413 | 0 | 0 | 0 | 0 | 0 | 0 | 0 | 1261 | 0 | 0 | 0 | possibly Australia | north Atlantic, north Pacific, South Atlantic, Hawaii, Mediterranean |
| Polychaeta | Heteromastus filiformis | 2 | 0 | 0 | 0 | 0 | 0 | 0 | 62 | 123 | 0 | 47 | 0 | 0 | 0 | 0 | 0 | 0 | 0 | 0 | 0 | 0 | cosmopolitan species complex | northeast Pacific |
| Polychaeta | Hydroides dirampha | 3 | 35 | 61 | 0 | 0 | 0 | 0 | 0 | 0 | 0 | 0 | 0 | 0 | 0 | 0 | 0 | 0 | 0 | 0 | 0 | 0 | possibly tropical western Atlantic or IndoPa | west Atlantic, northeast Pacific, Hawaii, Australia, New Zealand, Japan, Mediterranean |
| Scyphozoa | Blackfordia virginica | 1 | 0 | 0 | 0 | 0 | 0 | 0 | 0 | 0 | 119 | 0 | 0 | 0 | 0 | 0 | 0 | 0 | 0 | 0 | 0 | 0 | Black Sea | Atlantic, Caspian Sea, Pacific, southwest Atlantic |

| KEY | | | | |
|---|---|---|---|---|
| | reported native port | non-reported native | reported non-native | non-reported non-native |


REFERENCES

Algaebase: https://www.algaebase.org/

CABI: Invasive Species Compendium, https://www.cabi.org/

Diatombase: https://www.diatombase.org/

Barros, Rodolfo C., Rosana M. da Rocha, and Marcio R. Pie. "Human-mediated global dispersion of Styela plicata (Tunicata, Ascidiacea)." Aquatic Invasions 4.1 (2009): 45-57.

Crocetta, Fabio, Lionello Paolo Tringali, Henk K. Mienis, and Argyro Zenetos. "Clementia papyracea (Gmelin, 1791)(Mollusca: Bivalvia: Veneridae): its established status in the Mediterranean Sea and the first record from Greece." Cahiers de Biologie Marine 57.3 (2016): 271-275.

GLANSIS: Great Lakes Aquatic Nonindigenous Information System, https://www.glerl.noaa.gov/glansis/

NEMESIS: National Exotic Maine and Estuarine Species Information System, https://invasions.si.edu/nemesis/

NOBANIS: European Network on Invasive Alien Species, https://www.nobanis.org/

OBIS: Ocean Biodiversity System, https://obis.org/

PEP: Phytoplankton Encyclopedea Project

Schwindt, Evangelina, James T. Carlton, José M. Orensanz, Fabrizio Scarabino, and Alejandro Bortolus. "Past and future of the marine bioinvasions along the Southwestern Atlantic." *Aquatic Invasions* 15.1 (2020).

WoRMS: World Register of Marine Species, http://www.marinespecies.org/
